# Abundance and trends of harbor seals (*Phoca vitulina richardii*) in Alaska, 1996–2023

**DOI:** 10.64898/2026.09.02.748977

**Authors:** Peter L. Boveng, Gavin M. Brady, Michael F. Cameron, Cynthia L. Christman, Shawn P. Dahle, Lisa M. Hiruki-Raring, John K. Jansen, Stacie M. Koslovsky, Josh M. London, Brett T. McClintock, Robert A. Montgomery, Erin E. Moreland, Erin L. Richmond, Michael A. Simpkins, Jay M. Ver Hoef, Skyla M. Walcott, David E. Withrow, Jamie N. Womble, Kymberly M. Yano, Heather L. Ziel

**Affiliations:** Marine Mammal Laboratory, Alaska Fisheries Science Center, National Marine Fisheries Service, National Oceanic and Atmospheric Administration, Seattle, Washington, USA; Cooperative Institute for Climate, Ocean, and Ecosystem Studies, University of Washington, Seattle, Washington, USA; School of Environmental and Forest Sciences, University of Washington, Seattle, USA; University of Cumbria, Carlisle, United Kingdom; Resource Evaluation and Assessment Division, Northeast Fisheries Science Center, National Oceanic and Atmospheric Administration, Woods Hole, Massachusetts, USA; Glacier Bay National Park and Preserve, Southeast Alaska Network, National Park Service, Juneau, Alaska, USA; Protected Species Division, Pacific Islands Fisheries Science Center, National Marine Fisheries Service, National Oceanic and Atmospheric Administration, Honolulu, Hawai’i, USA

## Abstract

Pacific harbor seals (*Phoca vitulina richardii*) were surveyed throughout their range along Alaska coasts over a 28-year period. Twelve management stocks, delineated on the basis of evidence for demographic independence, were monitored for abundance and trends. We employed a two-stage Bayesian hierarchical analysis that integrated aerial survey counts with satellite-linked bio-logger haul-out timelines to account for the proportion of seals in the water and not visible to survey observers and cameras. Overall, harbor seals were abundant (201,122 seals in 2023) but, during the span of our study the population (stock) trends varied. The first half of our study was characterised primarily by growth, with a peak abundance of approximately 225,000 seals in 2015. Since then trends have been variable, with some stocks (e.g., Bristol Bay) showing continued growth while others (e.g. Prince William Sound, South Kodiak) declined. The recent regional declines coincided with observed climate anomalies (e.g., marine heatwaves) and the rapid retreat of tidewater glaciers. Our results provide a basis for managing this species in Alaska, which is protected under the Marine Mammal Protection Act; a vital nutritional and cultural resource for Alaska Native communities; and an important sentinel of change in the Northeast Pacific marine ecosystem.

## Introduction

Pacific harbor seals (*Phoca vitulina richardii*) are one of the most common and widespread marine mammal species in Alaskan waters. They inhabit the coastal waters of Southeast Alaska, west through the Gulf of Alaska and Aleutian Islands, and north to the Pribilof Islands and Bristol Bay in the Bering Sea (Allen and Angliss 2010, Figure 1, S1.1, S1.2). They feed primarily on fishes in marine and estuarine waters, but also in rivers and freshwater lakes—in some cases more than 100 km from the sea. Additionally, they haul out on rocks, reefs, beaches, frozen rivers and lakes, and drifting ice calved from glaciers. Harbor seals are important as upper-trophic predators in the coastal marine ecosystem, as a nutritional and cultural resource for Alaska Native communities, and as one of many natural attractions that draw visitors and commerce to Alaska. Estimates of harbor seal abundance and trend are vital for sound conservation and management and for planning of human activities in the marine environments where harbor seals occur.

**Figure 1.**
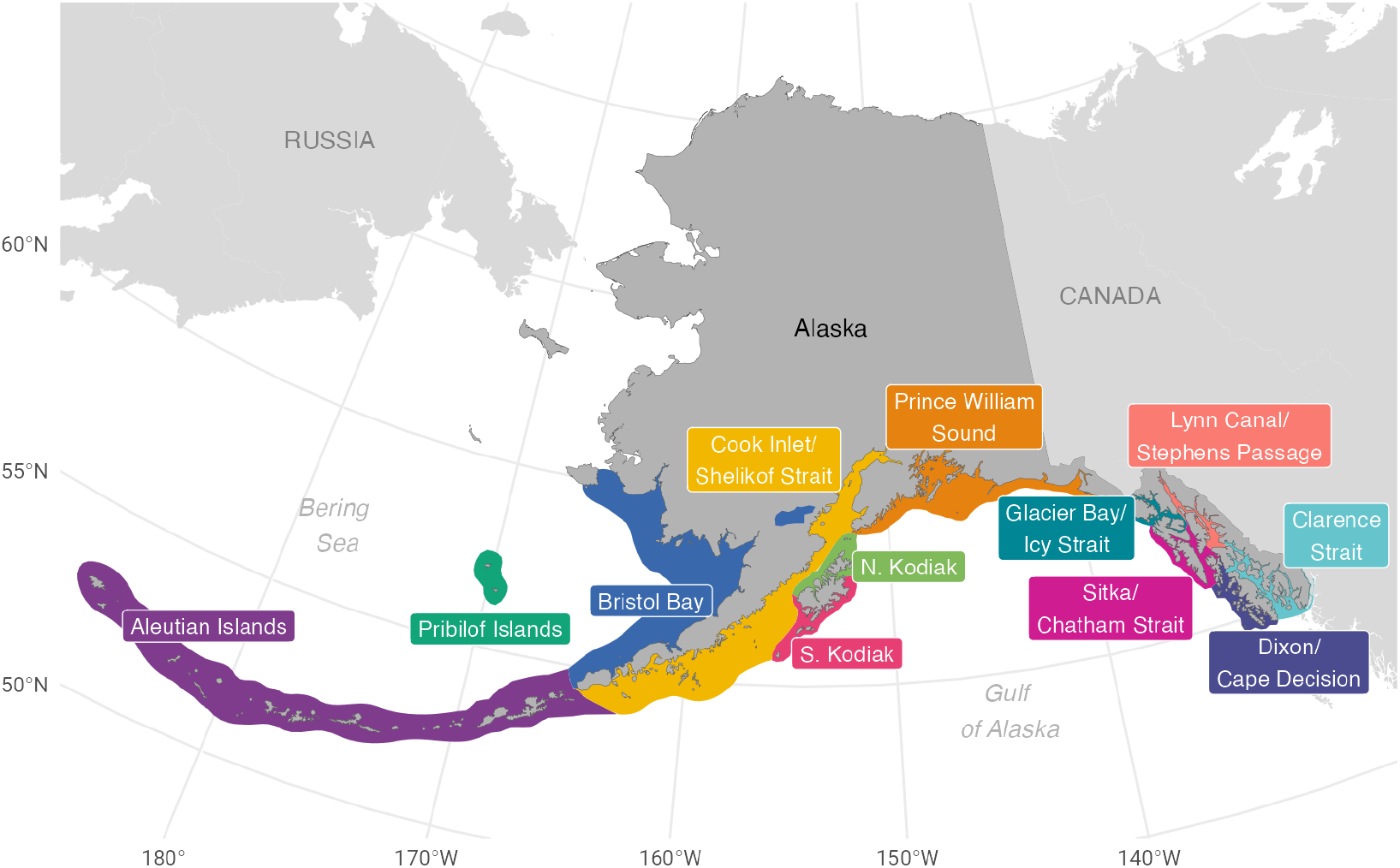
The geographic ranges and stock boundaries of harbor seals in Alaska.

Local and regional harbor seal numbers have been monitored at sporadic intervals since the 1970s, revealing various population trends. Where declines have been observed, they have been strongest in the late 1970s or early 1980s to the 1990s (Pitcher 1990, Frost et al. 1999, Ver Hoef and Frost 2003, Jemison et al. 2006, Small et al. 2008, Hoover-Miller et al. 2011, Womble et al. 2020), though estimates of harbor seal mortality from Indigenous subsistence, commercial, bounty, and predator-control harvests since the late 19th century imply that much larger declines likely occurred earlier (Crowell 2020). Some areas of more recent decline contrast with other regions where harbor seal numbers have remained stable or increased over the same period (Small et al. 2003), suggesting complex factors may be affecting population trajectories. In this study, we summarized and analyzed data gathered from a coordinated monitoring effort conducted throughout the harbor seal range in Alaska from 1996 to 2023, with the goal of documenting abundance and trends over the 28-year period.

Harbor seals can be surveyed most effectively when they are hauled out of the water and visible to survey observers or sensors. The numbers of harbor seals hauled out on shore or on ice vary seasonally and daily, influenced by many endogenous (physiological) and exogenous (environmental) factors; therefore, effective survey designs and reliable abundance and trend estimates rely on an understanding of haul-out timing. Harbor seals typically haul out in greatest numbers at lower tides (Pauli and Terhune 1987, Frost et al. 1999, Boveng et al. 2003, Sigourney et al. 2021, Ver Hoef et al. 2025) and during mid-day (Schneider and Payne 1983, Stewart 1984, Yochem et al. 1987, Watts 1996, Frost et al. 1999, Small et al. 2003, Mathews and Pendleton 2006), though these effects may vary by region (e.g. Frost et al. 2001, London et al. 2012, Boveng et al. 2018), by season (Cronin et al. 2010), or even by haul-out site (Stewart 1984, Ver Hoef and Frost 2003). There are two seasonal peaks in the numbers of harbor seals hauled out in Alaska: one during May/June associated with pupping and the other during August/September associated with molting (Jemison 2001). In Alaska, aerial surveys have mostly been conducted during the molting period when the number of seals hauled out is thought to be highest (Pitcher and Calkins 1979, Calambokidis and McLaughlin 1987) and the weather conditions are likely to be favorable for flying. Other environmental conditions such as local weather, substrate, and topography also affect the haul-out behavior of harbor seals.

Due to independent variation in tides, weather, and aerial survey logistical constraints, it is not possible to obtain counts at every harbor seal haul-out site under optimal conditions for seals to haul out. Thus, the inherent high variability in seal counts cannot be controlled by survey design alone, and the counts are typically adjusted to account for some or all of those factors. Such adjustments have been commonly used in trend analyses and abundance surveys of harbor seals in Alaska and British Columbia (Watts 1996, Frost et al. 1999, e.g. Boveng et al. 2003, Small et al. 2003, Ver Hoef and Frost 2003, Mathews and Pendleton 2006), as well as other regions (e.g., Sharples et al. 2009). In this study, we followed a similar approach and incorporated survey covariates into our analysis to reduce variability in estimates derived from harbor seal counts.

In addition to high variability, bias is a concern for harbor seal counts as estimators of total abundance because not all seals haul out of the water at the same time, even under ideal conditions of weather, tide, solar hour, and day of year. To estimate total abundance it is necessary to account for the fraction of the population that remains in the water during survey periods. We used satellite-linked bio-loggers to receive hourly timelines of the haul-out status from samples of harbor seals that were tagged in several regions throughout Alaska during the course of our study, which is the most common source of independent data used to account for the proportions of seals missed due to being in the water during abundance surveys (e.g., Sharples et al. 2009, London et al. 2024; Boveng et al. 2025).

The specific objectives of this study were to: (1) document a systematic monitoring framework for abundance and trends of harbor seal stocks in Alaska; (2) present a time series of abundance estimates and trends from 1996–2023 for each stock; (3) describe how environmental conditions affect the timing and numbers of seals hauled out during aerial surveys; and (4) consider our results in the contexts of harbor seal biology, management, and ecosystem dynamics.

## Methods

Over the duration of our study, our methods have consistently focused on aerial photographic surveys to obtain seal counts, the deployment of bio-loggers to collect haul-out behavior data, and the development of statistical methods to integrate these two data sources and relevant covariates into a robust estimate of abundance and trend. However, the multi-decadal duration of the survey effort means components of our protocols have adjusted, available technology has evolved, and personnel have changed.

Harbor seals in Alaska occupy two distinct habitat types: floating glacial ice and intertidal shores. Our approach differentiates aerial survey methods depending on whether the surveys are conducted over glacial fjords, where seals are dispersed in lower densities on floating ice, or over the coastline, where seals congregate in higher densities at intertidal haul-out sites. For surveys over glacial fjords, the methods have improved from visual counts and oblique photography (phased out by 2009) to gradual implementation of a GPS-integrated, downward-facing camera system (starting in 2002) to provide more accurate and consistent coverage of the drifting ice habitat (e.g., Jansen et al. 2015a, 2015b). For intertidal surveys, methods have consistently used handheld cameras to take oblique photographs of seals hauled out on varied substrates, such as rocky reefs, mudflats, and beaches. For images collected of both glacial and coastal habitats, seals were manually counted on photographic transparencies or digitally counted and/or mapped using image viewing or GIS software. Detailed aspects of the statistical framework underlying our estimates of abundance and trend are described in Ver Hoef et al. (2025). Here, we provide an overview of the main components of our long-term study. Those include: (1) designation of stock boundaries that are consistent with available genetics and movement data: (2) aerial survey design and allocation of effort across a large geographical area; (3) key methods and technologies for survey data acquisition and data management; (4) use of haul-out behavior data collected from bio-loggers; and (5) the application of advanced statistical methods towards a time series of estimates for abundance and trend.

### Harbor seal stock designation

In 2001, the Alaska Harbor Seal Co-management Committee—composed of harbor seal experts from the National Marine Fisheries Service (NMFS) and the tribally-authorized Alaska Native Harbor Seal Commission (ANHSC)—undertook an evaluation of the population (i.e., stock) structure of harbor seals in Alaskan waters (National Marine Fisheries Service 2002). The evaluation was based on genetic information, a geographic distribution of trends, movements of tagged seals, and traditional tribal hunting territories (in Southeast Alaska). Westlake and O’Corry-Crowe’s (2002) analysis of genetic information from 881 samples across 181 sites revealed harbor seal population subdivisions on a scale of 600–820 km, indicating that genetic differences within Alaska, and most likely over the entire North Pacific subspecies range, are positively correlated with geographic distance from each other. A subsequent analysis of smaller scale variation within the mitochondrial DNA (mtDNA) samples revealed substantial genetic differences indicating that female dispersal occurs at region-specific spatial scales of 150–540 km (O’Corry-Crowe et al. 2003). Although much of the geographic range of harbor seals in Alaska was unsampled, this research identified 12 clusters of sampling sites that were more likely to be demographically independent due to low dispersal rates between them.

In 2010, the NMFS and the ANHSC agreed to recognize 12 separate stocks of harbor seals based largely on the geographical distribution of mitochondrial genetic markers (O’Corry-Crowe et al. 2003). This was a marked increase in the number of harbor seal stocks from the three (i.e., Bering Sea, Gulf of Alaska, and Southeast Alaska) that were previously recognized. Because the genetic samples were not obtained continuously throughout the range, a ‘total evidence’ approach was used to consider additional factors, such as population trends (Frost et al. 1999, Small et al. 2003, 2008, Mathews and Pendleton 2006, Jemison et al. 2006), observed harbor seal movements (Lowry et al. 2001, Small et al. 2005, Blundell et al. 2011, Boveng et al. 2012), and boundaries of resource use areas traditionally controlled by Tlingit and Haida clans in Southeast Alaska (Goldschmidt and Haas 1998). These additional factors and the genetic results were displayed graphically and evaluated visually to determine boundaries by consensus among the NMFS and ANHSC co-management committee members. The boundaries were drawn to separate the 12 demographically-independent genetic sampling areas while—to the extent possible— separating areas of divergent population trends, avoiding the placement of boundaries across known harbor seal movements, and keeping traditional resource use areas within individual stocks of Southeast Alaska; the traditional resource use areas were noted by some committee members to be coincident with the genetic clusters in parts of that region.

The spatial areas represented in Figure 1 (see also Figures S1.1 and S1.2) were created to facilitate the analysis of abundance and trends and co-management of the 12 stocks of harbor seals in Alaska. The spatial boundaries extend up to approximately 70 km from the shoreline based on published movements of harbor seals (Lowry et al. 2001, Small et al. 2005, Boveng et al. 2012, Womble and Gende 2013, Cordes et al. 2017) and other geographic or oceanographic constraints. However, movements farther offshore do occur and the regions depicted in Figure 1 are not intended to define the full range of each stock. Also, the Bristol Bay stock includes a small resident population of harbor seals that lives within the fresh waters of Iliamna Lake, more than 115 km up the Kvichak River from Bristol Bay.

### Harbor seal aerial surveys

#### Survey design

As harbor seals can only be counted reliably when they are hauled out and visible on shore or on ice, our surveys covered the entire intertidal zone and all ice-covered areas of glacial fjords within the harbor seal range in Alaska (Figure 1). In earlier years, each seal count was associated with a general haul-out area where groups of seals were observed, typically what could be covered in one to a few photographs – which were marked on paper nautical charts and in later years recorded with GPS coordinates; these “waypoints” were refined post-survey so that they marked actual haul-out locations instead of the location of the survey aircraft. With real-time GPS mapping (i.e., moving maps), we recorded more precise positions and increased the granularity of haul-out locations, which led to increased variability in replicate counts due to the daily movements of seals among haul-out locations. Therefore, in 2006, we developed a standardized set of geographically distinct and identifiable seal survey units (SSUs) to serve as the smallest unit of record for aggregating seal counts (Montgomery et al. 2007, Boveng et al. 2011). Each SSU is an irregular polygon that follows the coastline and encompasses the intertidal zone, including offshore rocks or reefs. SSUs were delineated to be small enough (typically 12–15 km in length) to survey by airplane without significant changes in weather and tidal conditions (i.e., within 15–30 minutes) but large enough to avoid variations in counts from day-to-day differences in haul-out site preferences of local groups of seals. Seal counts and survey effort prior to 2006 were retroactively assigned to SSUs based on recorded haul-out locations, flight tracks, and observer notes.

Given the large geographic range of harbor seals in Alaska; challenging logistics and weather; inconsistent availability of suitable aircraft; and limited resources, spatial allocation of survey effort required complex planning and prioritization. Prior to 2008, effort was rotated, annually, through five different regions (i.e., southern SE Alaska, northern SE Alaska, Gulf of Alaska, Bristol Bay, and Aleutian Islands), and survey effort within each region was intensive with most SSUs surveyed multiple times. In 2008, annual survey effort was reduced (e.g., fewer flight hours) and shifted toward broader geographic coverage of stocks with the emphasis on high-abundance SSUs. Because the standard deviation of harbor seal counts scales with the square root of the mean (Taylor, 1961), sites with high expected abundance were prioritized to optimize the coefficient of variation (CV) for the total estimate (Cochran, 1977; Gasaway et al., 1986). Additional prioritization of SSUs was based on emerging conservation and management issues or time elapsed since the last survey.

#### Data acquisition

Surveys of harbor seals were flown with crewed, fixed-wing aircraft at altitudes between 225 and 1,219 meters (approx. 750-4,000 feet), and at ground speeds between 90 and 120 knots. In most years of the study, aerial imagery of seals hauled out at intertidal sites was collected at altitudes between 225 and 305 meters with handheld, oblique photography—the objective being to photograph every seal seen from the plane for post-survey counting. From 1996– 1998, the survey methods also relied regularly on visual counts recorded by observers in the plane (Boveng et al. 2003); since 1999, observers only recorded visual counts of small numbers of seals when photographs were not possible. Prior to 2003, color transparencies were captured with analog single-lens reflex (SLR) film cameras, and since then our methods have evolved to use digital SLR cameras, paired with geospatial information logged at the time of image capture. Survey timing was centered around daytime low tide events (2-3 hours either side of low tide and absolute tide heights generally less than 0.6 m) and the molting season for Alaska harbor seals (August through mid-September). This timing coincided with when we expect the highest proportion of seals to be hauled out. An accurate log of survey effort was an important component to our methods since 2006 to ensure that our dataset had a complete account of SSUs surveyed on a given day and that SSUs with no seals present (and no photos taken) were properly included in the final dataset. For years prior to 2006, we assigned effort based on a retrospective examination of flight tracks, photo locations, and observer notes. Starting in 2006, our methods required observers to categorize the survey as ‘recon’ (all known waypoints and potential haul-out habitat between the waypoints in the SSU were surveyed), ‘full’ (all known waypoints in the SSU were surveyed), or ‘partial’ (one or more waypoints in the SSU were not surveyed, typically due to weather constraints) during the flight. In this analysis, counts associated with ‘partial’ survey effort were not used.

The methodology for surveys of seals in glacial habitats also evolved over time. Prior to 2004, visual estimation or oblique photography methods typical of intertidal surveys were used within glacial habitats; these methods were supplemented in Glacier Bay with shore-based counts (Mathews and Pendleton 2006, Womble et al. 2020). In 2004, for specific glacial sites (Icy Bay and Disenchantment Bay) we began using a single, belly-mounted, downward-facing camera system better suited for surveys of seals on ice dispersed over the larger areas of glacial fjord habitats; this method was implemented in Johns Hopkins Inlet (Glacier Bay) starting in 2007 (Womble et al. 2020). Over subsequent years, we improved the system and expanded its use across most of the larger glacial fjord habitats in Alaska. By 2010, the system evolved into a three-camera array that reduced survey time while increasing spatial coverage and image quality, largely eclipsing the use of oblique photography in glacial fjord habitats. Surveys since 2010 were flown at higher altitudes (305–1,219 m (1,000-4,000 feet), depending on the fjord) compared to our surveys in coastal habitats. Surveys were generally conducted in the afternoon (approx. 12:00–17:00 h local time) when seals typically haul out on ice in peak numbers (Hoover 1983, Calambokidis et al. 1987, Blundell and Pendleton 2015, Mathews et al. 2016); surveys prior to 2002 were often flown in conjunction with intertidal surveys and often occurred before noon. The camera systems were configured to capture either full-coverage, overlapping imagery at higher altitudes (610–1,219 m) for a complete count of seals, or to collect non-overlapping imagery at lower altitudes (305 m) to support density estimation (Ver Hoef and Jansen 2015, Womble et al. 2020). The full-coverage approach was focused on smaller fjords that could be imaged in their entirety at higher altitudes (with larger image footprints) while maintaining adequate resolution for counting seals. This survey method entailed either capturing the full width of a fjord encompassing the ice-covered areas, or targeting a band of ice occupied by the vast majority of seals at a site. For larger fjords, where non-overlapping imagery was collected, the overall area sampled represented 20–48% coverage of the haul-out area within the fjord. Starting in 2020, a six-camera array (consisting of three color sensors paired with three long wavelength infrared [IR] sensors) provided high resolution color imagery with simultaneous automatic mapping of thermal “hotspots”. Thermal hotspots were manually confirmed as seals in the color imagery. The smaller IR footprint reduced spatial coverage of these surveys by 10–15%, but streamlined the analytical process. Digital image resolution of the color cameras improved as camera models were upgraded, resulting in ground sampling distances (GSD, distance between the centers of adjacent pixels in units on the ground) ranging from 2–4 cm/pixel, which was more than sufficient to discriminate seals on a lighter ice background.

There were three notable exceptions to our survey methods for intertidal and glacial habitats. First, in the Pribilof Islands, small uncrewed aircraft systems (sUAS) were used, starting in 2019, to collect vertical photographs of seals hauled out along the shoreline. Second, between 2001–2005, high-altitude photogrammetry (610 – 1,463 meters, Aeromap US Inc.; Bengtson et al. 2007) was conducted at all tidewater glacial sites known to have seals in Alaska. Third, in 2019, we evaluated the perceptibility of seals from the aircraft using a nose-mounted forward-looking infrared (FLIR) camera system in the Aleutian Islands (Christman et al. 2022). In that study, SSUs were selected and surveyed based on our experimental design for estimating perceptibility. Both color photos and FLIR imagery were collected and counted but only counts from the color photos were used to estimate abundance. Counts from the FLIR images informed our estimate of perceptibility (see Abundance Model section below). In all situations, accurate time values and precise spatial coordinates were recorded for each image such that resulting counts could be accurately assigned to a SSU.

#### Data processing and management

The management of data collected during the aerial surveys evolved in concert with the technology and types of data acquired. For surveys in all habitats from 1996–2002, color transparencies from analog cameras were projected onto a white background, and the seals were counted. This was typically completed by two independent counters and the mean of the two counts was used. For all images taken since 2003, the Exchangeable image file format (Exif) metadata associated with images from digital cameras was extracted from the images and imported into a relational database. This captured key information including date, time, and geospatial coordinates (either collected directly from a GPS attached to the camera or via a post-hoc analysis of the flight track). Digital images were examined, and seals were counted within image analysis (e.g., Adobe Photoshop, Kitware’s DIVE) or GIS software (Esri ArcGIS or QGIS). Prior to 2015, for full-coverage glacial surveys, image mosaics were assembled with commercial imaging and GIS software to acquire full counts without missing or double-counting seals. Since 2015, we have increasingly employed advanced scripting and custom R software (R Core Team 2023) to streamline the process of preparing images for counting. Counts for high-altitude imagery were done visually; for low-altitude imagery, we used a detection model (YOLOv3, Redmon and Farhadi 2018; see also https://doi.org/10.5281/zenodo.5765664) to detect IR hotspots, with visual review of the corresponding color imagery to confirm the presence of each seal.

Upon completion of the surveys, the information from field datasheets was manually entered and the GPS tracklines were imported into a PostgreSQL relational database (http://www.postgresql.org/) with the PostGIS extension (http://postgis.net/); entries were reviewed and queried to assure the quality and accuracy of the data. Tidal covariates were determined for each SSU surveyed based on the most relevant (i.e., nearest by water) tide station and the date and time of the survey. Tidal covariates were predicted via the command line interface for XTide (https://flaterco.com/xtide) with updated harmonics parameters. Custom functions within R were developed to estimate time to the nearest low tide, tide height at the time of survey, and tide height for the nearest low tide and nearest high tide.

### Haul-out timelines

Satellite-linked bio-loggers were deployed on free-ranging harbor seals by the Alaska Department of Fish and Game (ADF&G) (Womble et al. 2020), the NMFS, and the National Park Service (NPS) between 2004 and 2017 (Womble and Gende 2013). Some details of the capture, tagging, and release methods varied with the study objectives and locations, but all bio-loggers relied on a conductivity sensor to determine the wet/dry status of the tag. Those observed values were summarized into hourly percent-dry timelines onboard for transmission efficiency and, later converted to binary values (above or below 50%) for input into our analysis (see London et al. 2024 for additional details).

### Analysis

To estimate population abundance and trends for each stock, we combined separate analyses of the haul-out data and the aerial survey data using a hierarchical approach. Complete details of this two-stage Bayesian analysis (including prior specification and alternative parameterizations) can be found in Ver Hoef et al. (2025), but we provide a general overview of the methods here.

The survey count data are informative about population abundance, but they are subject to imperfect detection because only seals that are hauled out during surveys are ‘available’ to be counted (i.e., availability bias; Marsh and Sinclair 1989). The counts are therefore intricately linked to the proportion of seals that are hauled out when the survey aircraft passes. We accounted for this in our two-stage approach by first fitting an availability model based on the haul-out data and then incorporating this information into an abundance model based on the survey count data. All models were fitted using a custom-coded Markov chain Monte Carlo (MCMC) algorithm (See Appendix A in Ver Hoef et al. 2025) in R (R Core Team 2023). While our MCMC algorithm was very similar to that of Ver Hoef et al. (2025), we made several key changes as detailed in Supplemental Material S3.

### Availability model

In the first stage of the analysis, the (binary) haul-out data (ℎ_*t*_ ∈ {0,1}) were modeled using a logistic regression with temporally-autocorrelated (ϵ_*t*_) and individual-level (*γ*) random effects. Ignoring individual-level subscripts for simplicity, we have:

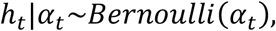

where the haul-out (i.e., availability) probability at time *t* can be expressed as:

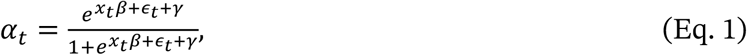

where *x* = (1, *d, d*^2^, *u, u*^2^, *l, l*^2^) is a vector of explanatory variables, d ∈ {−30,45} is day of year from 15 August, *u* ∈ {−12,12} is hours from solar noon, *l* ∈ {−5,5} is hours from low tide, and β is the corresponding vector of regression coefficients. Quadratic terms were included because we expected each variable to have a peak (e.g., around 15 August, solar noon, and low tide, respectively). Whereas the haul-out data were typically spaced 1 hour apart, the temporally-autocorrelated random effects (ϵ_*t*_) were specified such that they could accommodate irregularly-spaced observations, such as when there were missing data. For numerical stability and efficiency, all random effects were modeled using truncated normal errors (see Ver Hoef et al. 2025).

Noting that haul-out data were sparse or absent for some stocks, and suspecting that haul-out behavior would likely differ based on geographic location and intertidal versus glacial ice haul-out substrates, we fit four separate haul-out models. Three models were fit to haul-out data most closely linked geographically to the intertidal SSUs in three groups of stocks (Table 1). The fourth model was fit to haul-out data from the glacial fjords where harbor seals haul out on ice in the Prince William Sound, Glacier Bay/Icy Strait, Lynn Canal/Stephens Passage, and Clarence Strait stocks. For the glacial haul-out model, we excluded hours from low tide.

**Table 1.** Sample sizes (hourly values of haul-out status) and numbers of individual seals with bio-logger deployments for fitting availability (i.e., haul-out) models. Haul-out records from all glacial SSUs were combined and modeled separately from SSUs on shore in the stocks where the glacial fjords are located.

| Group | Stocks | Age | Sex | Number of Records | Number of Seals |
| --- | --- | --- | --- | --- | --- |
| Western Islands | Aleutian Islands,<br>Pribilof Islands | Young-of-Year: 11<br>Subadult: 25<br>Adult: 39 | Female: 40<br>Male: 35 | 28,370 | 75 |
| Southwest and Central Alaska | Bristol Bay, Cook Inlet/Shelikof Strait, South Kodiak, North Kodiak, Prince William Sound | Young-of-Year: 10<br>Subadult: 10<br>Adult: 26 | Female: 25<br>Male: 21 | 16,007 | 46 |
| Southeast Alaska | Glacier Bay/Icy Strait, Sitka/Chatham Strait, Lynn Canal/Stephens Passage, Dixon/Cape Decision, Clarence Strait | Subadult: 8<br>Adult: 25 | Female: 24<br>Male: 9 | 21,342 | 33 |
| Glacial Fjords | Glacial SSUs in Central and Southeast Alaska | Subadult: 6<br>Adult: 19 | Female: 18<br>Male: 7 | 9,463 | 25 |

### Abundance model

In the second stage of the analysis, we estimated abundance for SSU *j* and year *t* B*N*_j,*t*_D of each stock based on the aerial survey counts and the (first-stage) availability models. As detailed in Ver Hoef et al. (2025), we modeled the *i*th count from SSU *j* in year *t* B*c*_*i,j,t*_D using a binomial distribution:

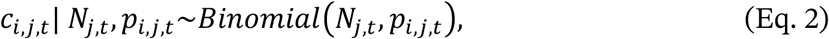

where

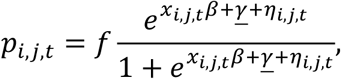

and *f* is the probability of perception (‘perceptibility’) for a seal that is hauled out, *x*_*i,j,t*_ is the vector of explanatory variables associated with *c*_*i,j,t*_, *β* is the vector of corresponding coefficients, *γ* is the average of the individual-level effects from the haul-out model (Eq. 1) corresponding to the stock, and *η*_*i,j,t*_ is a random effect to allow for overdispersion for each survey. Notably, the priors for *β* and *γ* were obtained from the joint posterior distribution of *β* and *γ* from the haul-out model for the stock. While for most stocks we assumed *f* = 1 (i.e., all individuals that are hauled out and within a SSU are counted), there was concern about reduced sightability for seals in the Aleutian Islands stock stemming from our observations of generally smaller group sizes and darker pelage (Shaughnessy and Fay 1977) that more easily blends in with the typically dark, rocky haul-out areas in the Aleutian Islands. For this stock, we computed the posterior distribution of *f* using data from a study that used FLIR cameras (Christman et al. 2022, Ver Hoef et al. 2025) as an independent detection method. We used a simple Bayesian Poisson regression model to obtain the posterior distribution of this fraction, which had a mean of *f* = 0.86. We used the posterior distribution of *f* as a prior distribution in our model for counts in the Aleutian Islands stock.

Due to challenges associated with surveying some of the larger glacial fjords, an additional complication with these particular surveys is that they do not constitute complete counts (i.e., coverage). Rather, the complete counts for these glacial SSUs were estimated (along with an associated estimate of variance) by Ver Hoef & Jansen (2015). As detailed in Ver Hoef et al. (2025), the estimated counts for these glacial surveys were modeled using a Beta-Binomial distribution (which is parameterized such that it reduces to Eq. 2 when there is no uncertainty in the count).

For the SSU-level population size at time *t*, we assume:

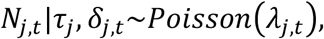

where *λ*_j,*t*_ =*exp* Bτ_j_ + δ_j,*t*_D, τ_j_ is the intercept for SSU *j*, and δ_j,*t*_ is a temporally-autocorrelated random effect that is assumed to follow a random walk (see Ver Hoef et al. 2025 for further details). There are several important properties of the random walk in this formulation. First, in the absence of any further counts, the expectation of δ_j,*t*+*k*_ |δ_j,*t*_ is δ_j,*t*_regardless of *k* (the temporal projection horizon), which means that δ_j,*t*_ projected into the future (or past) will tend to remain constant after the last year in which surveys were conducted. Second, in the absence of any further counts, the variance of δ_j,*t*+*k*_ |δ_j,*t*_ grows linearly without bound as *kσ*_δ_^2^ where *σ*_δ_^2^ is the variance in the change in δ_j,*t*_ from year to year. Finally, δ_j,*t*_ is modeled on the log scale, and upon exponentiating it has a lognormal distribution. The expectation of exp(δ_j,*t*+*k*_ |δ_j,*t*_) is exp(δ_j,*t*_ + *kσ*^2^ /2), so that both the mean and the variance of the posterior increase with *k*. Thus, in the absence of any further counts after time *t*, N_j,*t*+*k*_ will tend to drift upwards because the random walk is on the log scale and *N*_j,*t*_ is bounded at zero. To mitigate this upward drift, we therefore included the weakly informative prior *λ*_j,*t*_| *α*_*λ*_, *β*_*λ*_*∼Gamma(α*_*λ*_, *β*_*λ*_), with shape parameter *α*_*λ*_ = 1 and rate parameter *β*_*λ*_ = 0.001. More details of consequences of these effects are given in the Results section.

The survey counts from within the stock were modeled jointly:

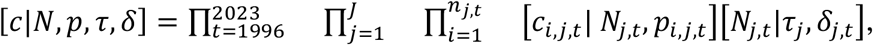

where *J* is the number of SSUs in the stock and *n*_j,*t*_ is the number of replicate counts at SSU *j* in year *t*.

Total abundance and trends were computed from the posterior distributions of *N*_j,*t*_ by summing across all *j* within a stock for year *t*. Because we had samples of the posterior distributions of all *N*_j,*t*_, we also had samples from the posterior distribution of any sum of these, which was the basis for credible intervals. Trends were computed by regressing the yearly abundance estimates on the trailing 8 years, providing an estimate of average yearly change in abundance during the 8 years leading up to the current one. A similar regression on the log-transformed abundances, produced an average percent change per year over the same time span. The 8-year span was chosen as a balance between shorter spans that have lower statistical power to detect trends and longer spans that provide a less current assessment of the trend. An 8-year span also has figured prominently in the evolution of NMFS guidelines for the assessment of marine mammal stocks, particularly for evaluating the reliability of abundance estimates as they age (Wade and Angliss 1997, National Marine Fisheries Service 2005, 2016, Moore and Merrick 2011, Bettridge 2023). In a similar fashion to obtaining samples from the posterior distribution of abundance, we obtained the samples from the posterior distribution of trends (in absolute yearly change, or percent yearly change) and computed their credible intervals (see Ver Hoef et al. 2025 for further details).

A different analysis was used to estimate the abundance of harbor seals resident year round in Iliamna Lake, which compose approximately 1.5% of the Bristol Bay stock. Aerial surveys for seals in Iliamna Lake were conducted from 1984 through 2024 (Withrow et al. 2015 and this study). This effort often included multiple surveys per year across different seasons and, during pupping season, some surveys distinguished pups from non-pups. These data were more conducive to a Bayesian integrated population model that combined counts—including pup counts—with a simple age-structured population model (Boveng et al. 2018) to produce a time series of yearly estimates for the abundance of seals in Iliamna Lake. These yearly posterior samples were simply added to those from our main model, yielding posteriors for the total Bristol Bay stock abundance.

## Results

### Spatial and temporal extent of surveys

Our surveys, conducted between July and September of 1996–2023, comprised 872 flights totaling more than 3,000 hours by observers in aircraft; there were also 45 flights of sUAS to survey the Pribilof Islands stock. The entire range over which harbor seals were observed to haul out on Alaska shores and floating ice comprised 1,272 SSUs and covered roughly 4,000 km from Southeast Alaska to the western Aleutians and 53 deg of longitude. The mean and median number of counts at each of the SSUs were 17 and 9, respectively.

### Alaska-wide abundance and trends

We estimated the combined abundance from all 12 stocks of harbor seals in Alaska in the summer (i.e., molting period) of 2023 to be 201,122 individuals, with a coefficient of variation of 0.018 and a 95% credible interval of 194,459–208,174. Our retrospective estimates of Alaska statewide abundance ranged from approximately 180,000 in the late 1990s to a high of about 225,000 in 2015, then declined to around 200,000 through the early 2020s (Figure 2). The annual estimates of the trailing 8-year trend were positive from 2003– 2016, with a maximum annual increase of 2.04% in 2006. In the years since 2016, the annual estimates of trend have been negative with a maximum decline of-1.71% in 2021 (Figure 2).

**Figure 2.**
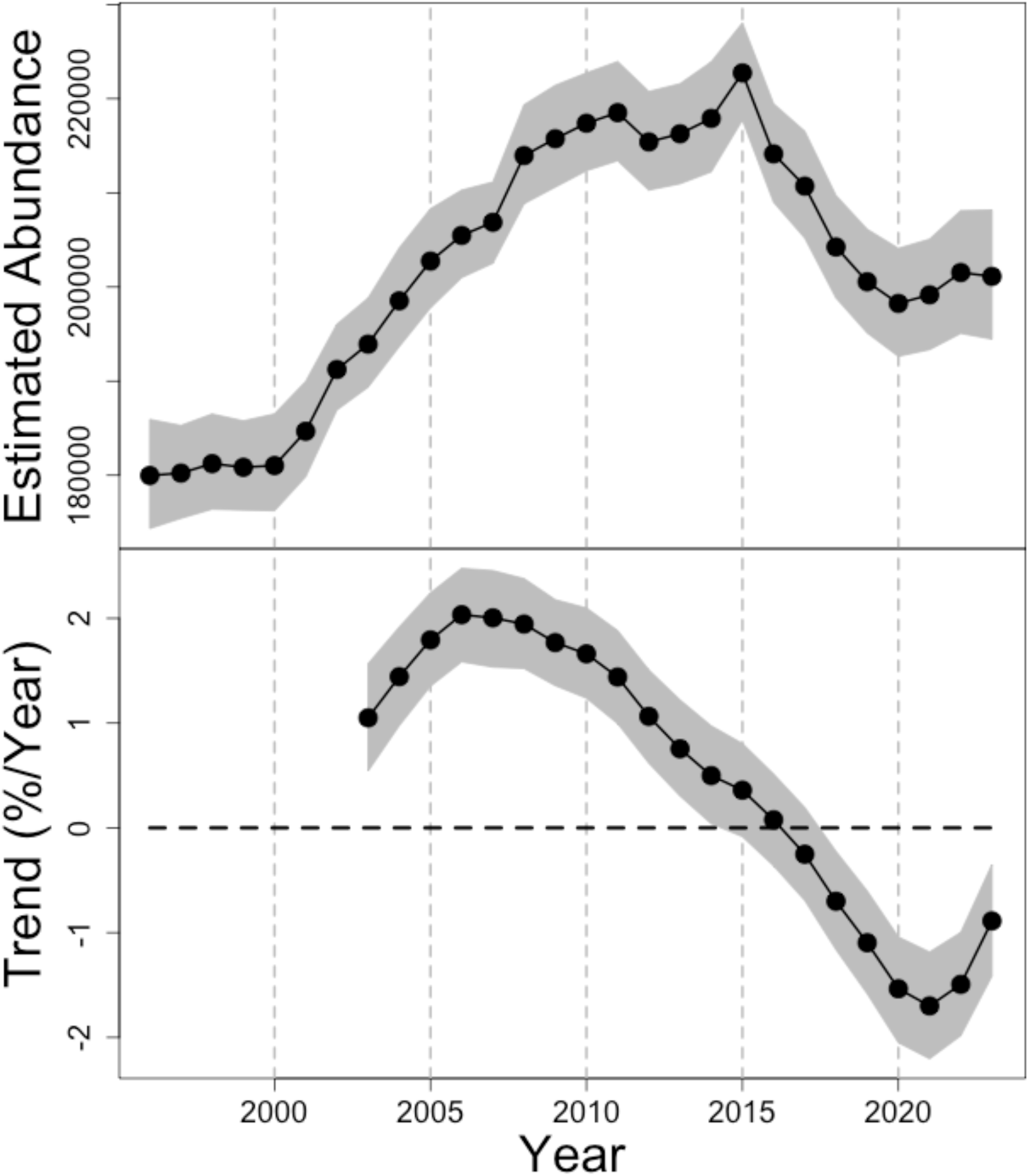
Estimated total number of harbor seals in Alaska from 1996–2023 (black circles, upper panel) and the annual trailing 8-year trend estimates, expressed as %/year (lower panel). The gray-shaded regions represent 95% credible intervals for the estimates.

### Variation among stocks

#### Abundance estimates in 2023

The 12 harbor seal stocks differed substantially in their abundances. The 2023 estimates (Table 2) ranged from 37,142 individuals in Bristol Bay, the largest stock, to 355 individuals in the Pribilof Islands, the smallest. Four stocks—Bristol Bay, Cook Inlet/Shelikof Strait, Prince William Sound, and Clarence Strait—represent 60.75% of the total statewide abundance.

**Table 2.**
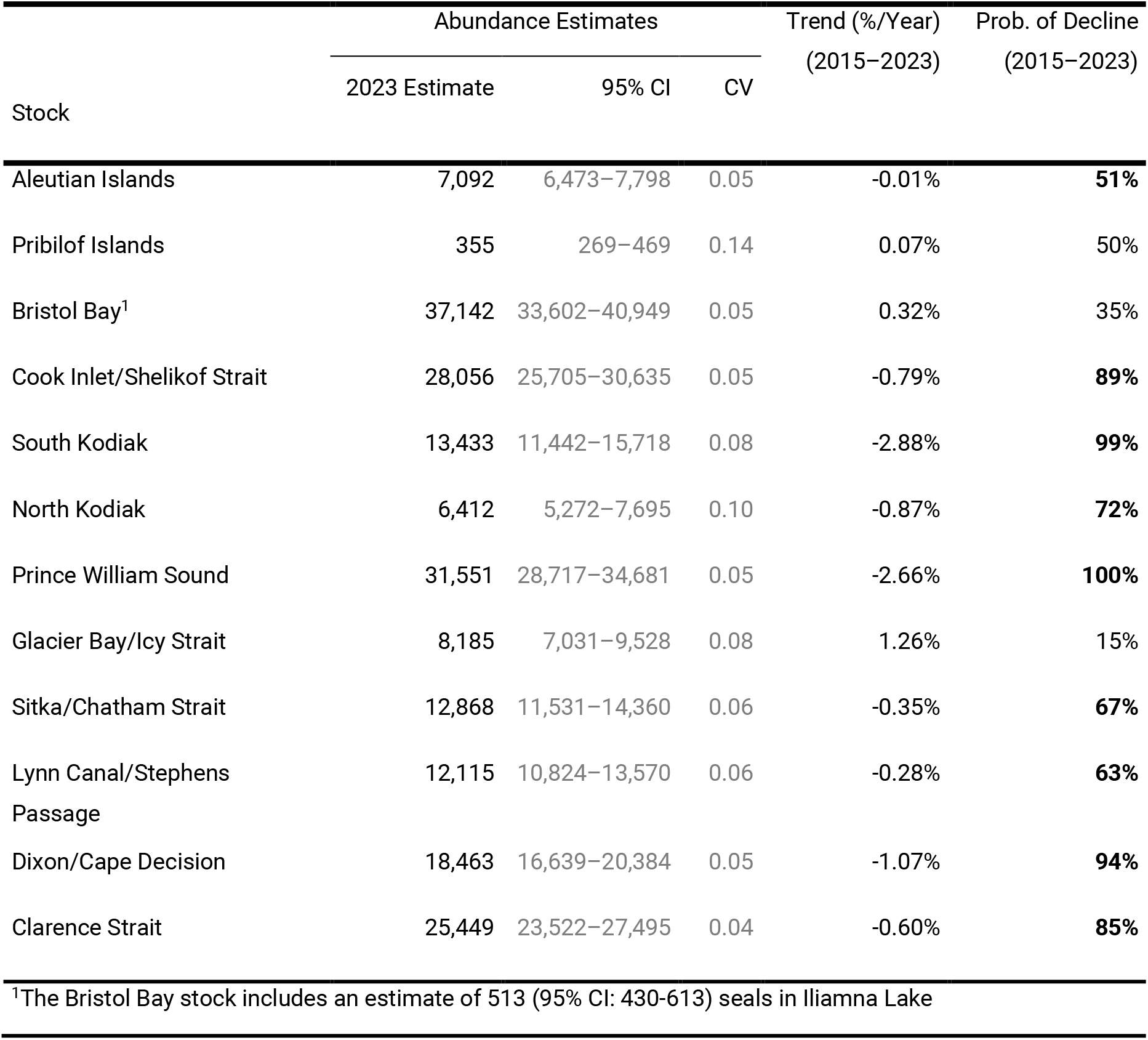
Abundance and trend estimates in 2023, by stock, for harbor seals in Alaska, with respective estimates of precision (i.e., the 95% credible interval [CI] and coefficient of variation [CV]). The trend estimate represents the annualized percent change in abundance from 2015 to 2023, as an index of current trend. The probability of decline represents the proportion of the posterior distribution of the trend estimate that is less than 0. Stocks more likely to be declining (*p*>50%) are highlighted with bold font. Due to rounding, the sum of abundance estimates across all stocks may not equal the reported total abundance of 201,122.

#### Population trajectories (1996–2023)

The estimated annual abundance and associated 95% credible interval for each year from 1996–2023 (Figure 3) provide a visual basis for examining the long-term patterns of growth and decline for each of the 12 harbor seal stocks in Alaska. In the lower panel of each subfigure, we provide an annual measure of survey effort expressed as the sum of abundance estimates across SSUs surveyed in that year divided by the stock-wide abundance estimate. This is an estimate of the proportion of seals in the stock surveyed in a given year. To illustrate with a simple example, suppose a stock was composed of 5 SSUs with estimated abundances in a particular year of 10, 2, 100, 50, and 80 seals; suppose that the 1st, 3rd, and 5th SSUs were surveyed in that year. In this example, the effort would be (10 + 100 + 80)/(10 + 2 + 100 + 50 + 80) = 0.79; in Figure 3, this effort index is shown as a gray bar for each year.

**Figure 3.**
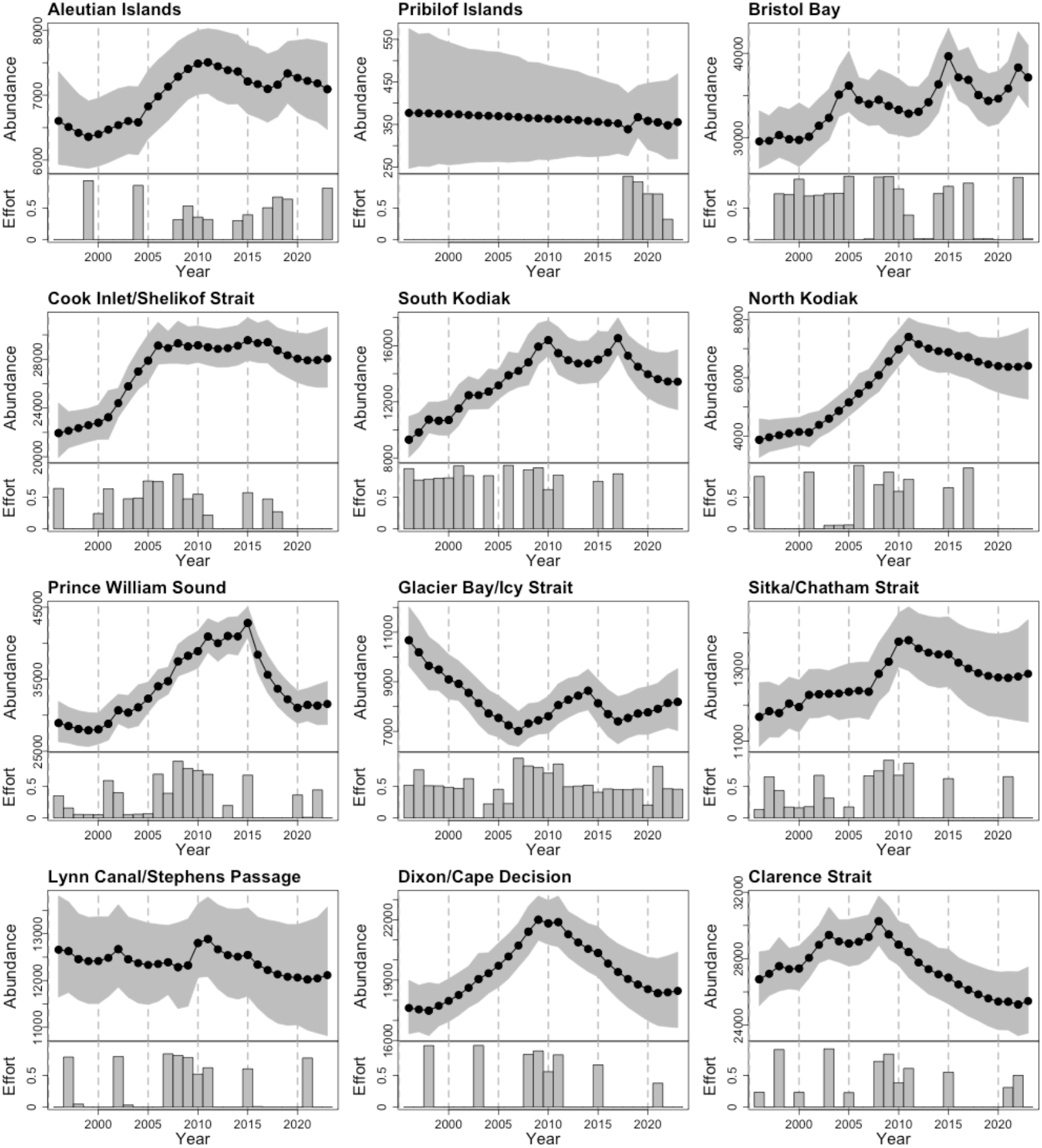
Abundance estimates (black circles) and survey effort (gray bars) for the 12 stocks of harbor seals in Alaska, 1996–2023. Ninety-five percent credible intervals are displayed as gray-shaded regions in the upper panel of each subfigure. In the lower panel of each subfigure, our index of survey effort was calculated as the sum of the estimated abundance for those SSUs that were surveyed at least once in that year, divided by that year’s estimated abundance for the entire stock.

As mentioned in the Methods section above (further detail in S3), preliminary analyses indicated that the SSU-level estimates of abundance have a tendency to drift upward over time in the absence of subsequent counts. This non-significant trend was especially evident when there were several years without counts at the end of a time series (e.g., Cook Inlet/Shelikof Strait, South Kodiak, North Kodiak). To counteract this tendency from our model structure, we included a weakly informative Gamma prior. The estimates presented in Figure 3 provide a more sensible projection forward in the absence of counts, with the point estimates remaining relatively stable and the credible intervals expanding over time.

#### Population trends

An estimate of the current rate of population change is a fundamental component of stock assessment and effective population management. Our statistic for the current rate of population change, the 8-year trailing trend in 2023 expressed as percent change per year (Table 2), was significantly different from zero for two stocks. In 2023, the Prince William Sound stock trend estimate was-2.66%/year and the South Kodiak stock was-2.88%/year with corresponding posterior probabilities of decline of 1.00 and 0.99. The probability of decline in 2023 is the proportion of the posterior distribution for the trailing 8-year trend that lies below 0. Among the remaining stocks, the probability of decline ranged from 0.15 for Glacier Bay to 0.94 for Dixon/Cape Decision. For 9 of the 12 stocks, the probability of decline was greater than 0.5. Among the hindcasted 8-year trend estimates for previous years (Figure 4), all stocks except the Pribilof Islands and Lynn Canal/Stephens Passage had periods of significant growth and/or decline during the monitoring period.

**Figure 4.**
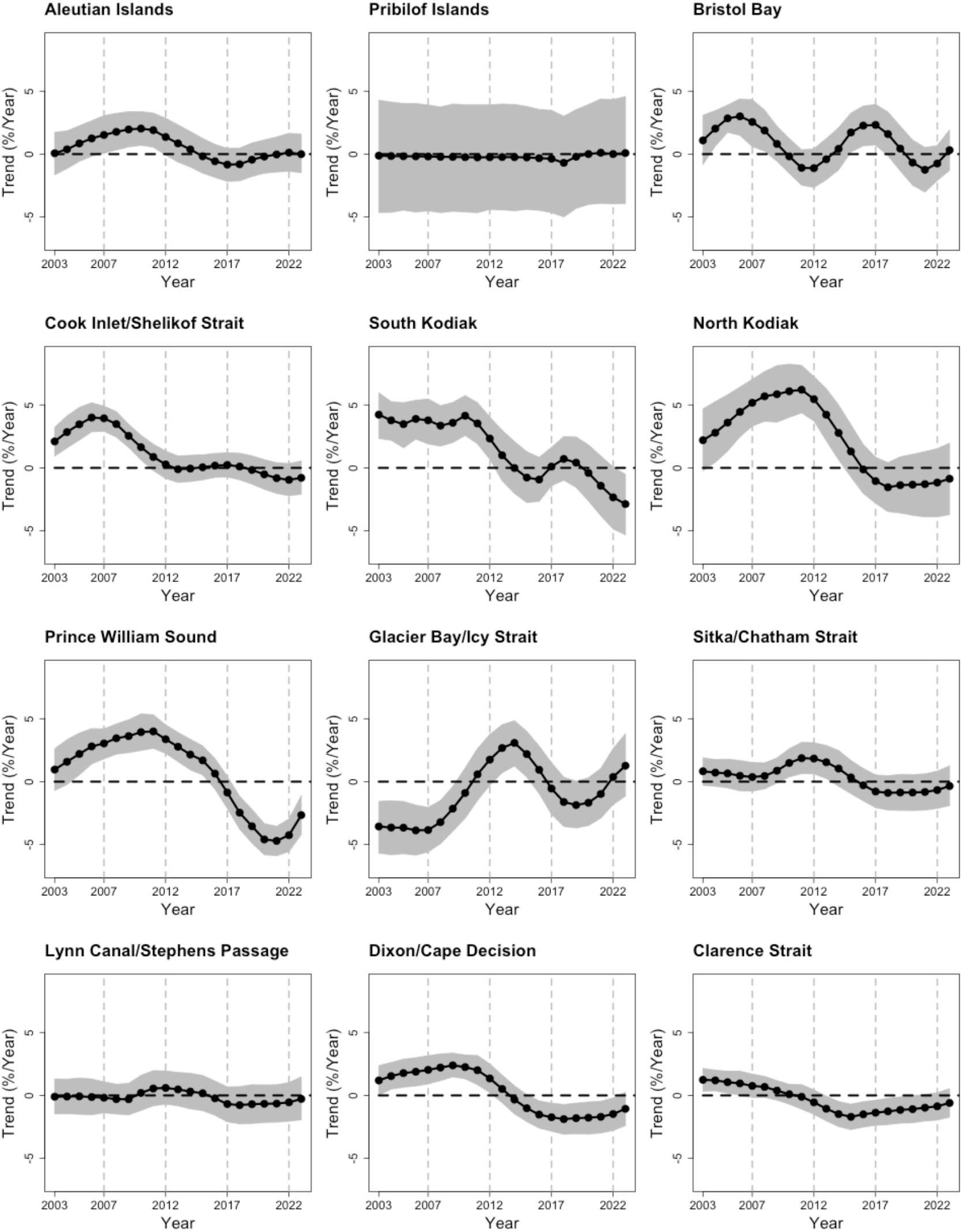
Time series of 8-year trailing trends for each harbor seal stock during 2003–2023, expressed as percent change per year (black circles). The gray-shaded regions are 95% credible intervals for the trend estimates.

Although the 8-year trend estimates are the best indicators of recent population change, there may also be interest in long-term change during the monitoring period. To gain insight into whether there may be regional patterns in the long-term trends among stocks, we arranged simplified depictions of the abundance time series in a map orientation (Figure 5). We consider these patterns as potential indicators for ecosystem dynamics in the Discussion.

**Figure 5.**
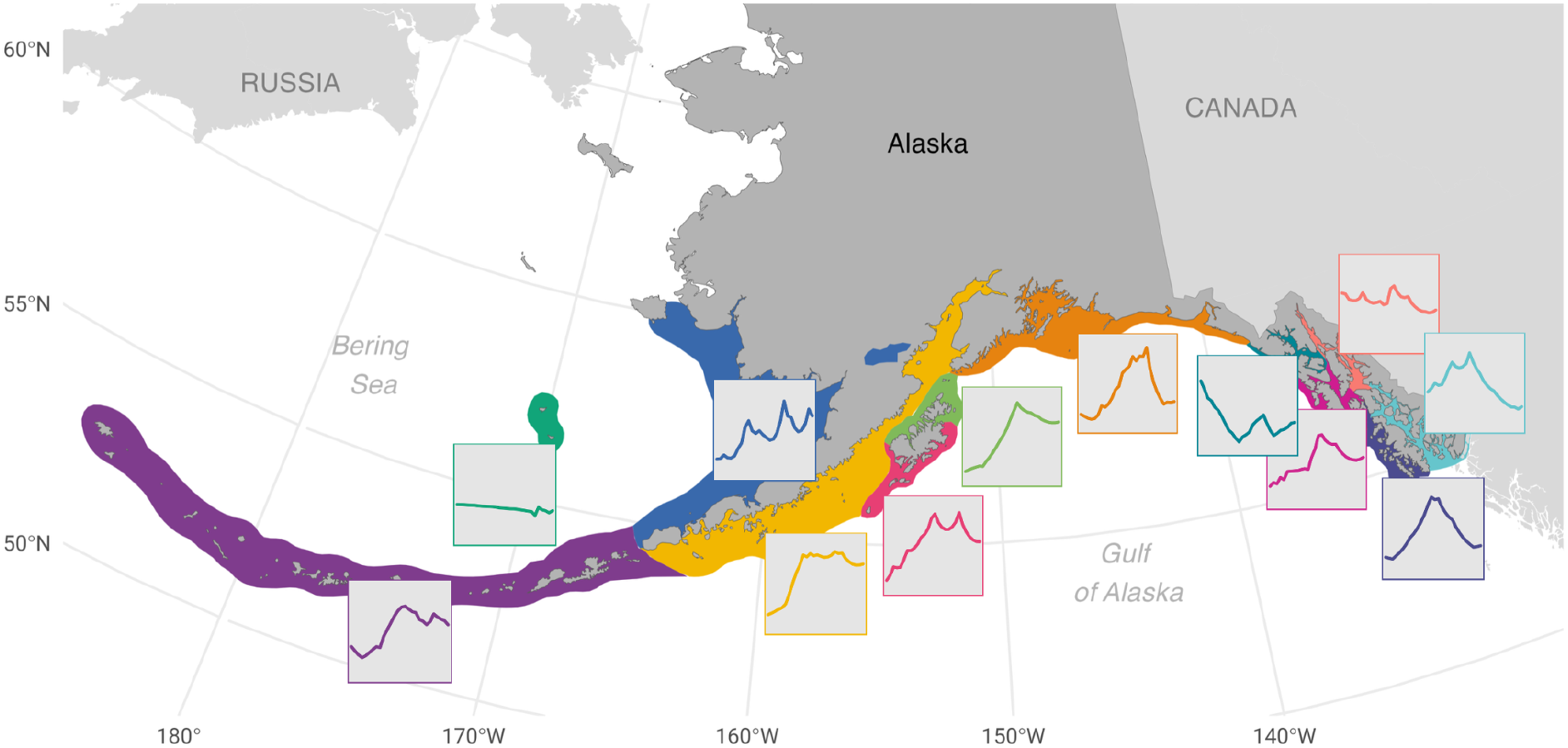
Map-oriented view of abundance time series for the 12 stocks of harbor seals in Alaska, 1996–2023. Stock boundaries are presented in the same colors as Figure 1. Stock names in order from east to west: Aleutian Islands, Pribilof Islands, Bristol Bay, Cook Inlet/Shelikoff, South Kodiak, North Kodiak, Prince William Sound, Glacier Bay/Icy Strait, Sitka/Chatham Strait, Lynn Canal/Stephens Passage, and Dixon/Cape Decision.

### Explanatory variables

The influences of date, hours from solar noon, and hours from the nearest low tide can be seen in Figure 6 for the three regional groupings of haul-out data from seals hauling out ashore and the grouping for glacial SSUs (Table 1), where the hour from low tide has no effect on availability of haul-out space (i.e., floating ice).

**Figure 6.**
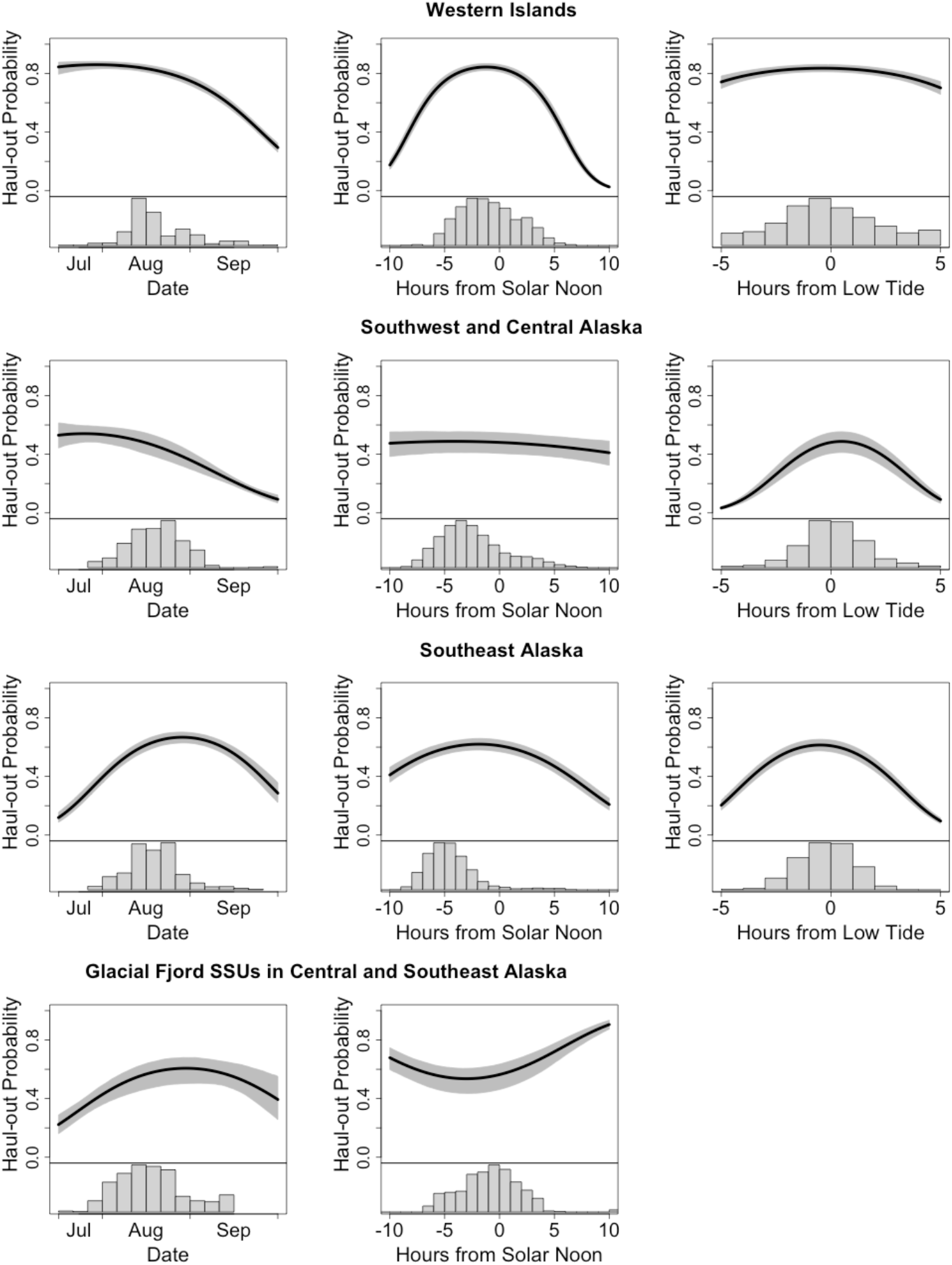
The effects of date, hour of day, and hours from low tide on haul-out probability for three groupings (Table 1) of haul-out records from harbor seals hauling out ashore (top three rows). The bottom row depicts effects of date and hour of day in haul-out records from glacial fjords, where the tide does not influence availability of floating ice for hauling out (Boveng et al. 2003). The groupings indicate the stocks (or glacial SSUs) to which the haul-out probabilities were applied; no haul-out records were available from the Pribilof Islands and Lynn Canal/Stephens Passage stocks. Histograms below the fitted curves show relative distributions of aerial survey data in relation to the fitted curves.

## Discussion

Along with a detailed description of our statistical methods (Ver Hoef et al. 2025), we have documented a comprehensive summary of our efforts monitoring harbor seal abundance and trends in Alaska over a 28-year period. The results presented here represent the current, best estimates of abundance and trend for harbor seals in Alaska between 1996 and 2023 and should supplant any previous publication of estimates derived from subsets of our survey and monitoring data (e.g., NMFS Marine Mammal Stock Assessment Reports). This serves as foundational information on abundance and trends required for assessment and management of harbor seal stocks under the U.S. Marine Mammal Protection Act (MMPA; Marine Mammal Commission 1995, NOAA 2023, NOAA Fisheries 2023).

### Monitoring, assessment, and management

Our analysis yields posterior probability distributions for abundance and trend estimates. They inherently account for uncertainty from many sources and so are well-suited for stock assessments. For example, *N_min_* —the minimum population estimate that is used under the MMPA to determine the potential biological removal (PBR; the allowable level of human-caused mortality and serious injury)—is based on the 20th percentile of the probability distribution for the abundance estimate (Barlow et al. 1995, Wade 1998). The 20th percentile can be extracted directly from our posterior distribution for abundance, rather than using an estimate that relies on assumptions about the shape (e.g., lognormal) of the probability distribution for an estimator (e.g., Wade 1998).

Another product of our model framework, the probability of decline, has utility for determining the recovery factor, *F_r_*, (Barlow et al. 1995, Wade 1998) in the PBR framework. *F_r_* ranges from 0.1 to 1.0, and serves to reserve a portion (i.e., 1−*F_r_*) of the population’s net production for growth rather than allowable take. *F_r_* also provides a means of compensating for uncertainty or bias in other parameters or assessment data that might impede population recovery. In practice, *F_r_* is determined partially on the basis of judgment by federal agency managers and an independent scientific review group, applying criteria such as whether the stock is depleted, threatened, of unknown status, or is within its optimum sustainable population range; whether it is increasing or declining; whether it is taken primarily by Indigenous subsistence hunters; and its recent history of mortality and serious injury, along with the precision of those estimates (Bettridge 2023). Our estimates for the 8-year trailing trend and the probability of decline provide direct, quantitative inputs to two of those criteria for consideration in the selection of *F_r_*. The 8-year trend estimate, as the average of the posterior distribution for the trend, provides the ‘best’ estimate of the rate of change in the population. The probability of decline, as the proportion of the posterior distribution that lies below zero, provides context as a measure of the likelihood that the population is actually declining.

### Ecosystem dynamics

Congruent dynamics among populations distributed over a large geographic area may reveal patterns driven by broad-scale ecological processes and, by similar logic, differences in dynamics of adjacent populations may be indicative of smaller-scale variation in such processes. For example, with the exception of the insufficiently monitored (until recently) Pribilof Islands stock, all but two stocks (Glacier Bay/Icy Strait and Lynn Canal/Stephens Passage) had net increases (positive linear trend value and 95% credible interval not spanning 0.0; see Table S2.1) in abundance through the first half of our monitoring period (i.e., 1996–2009; Figures 3 and 4). This prevalent pattern in the first half of the study period could reflect broad-scale drivers of population growth across much of the range of harbor seals in Alaska.

The results of monitoring (mostly) prior to this study may also suggest broad-scale coherence in population dynamics, though the prevalent pattern found was a decline in harbor seal numbers from the late 1970s or early 1980s into the 1990s or 2000s in some areas. Long-term counts at Tugidak Island (now in the South Kodiak stock) declined by more than 80% from 1976 through 2000 (Pitcher 1990, Jemison et al. 2006). Counts at Nanvak Bay in northern Bristol Bay declined by 84% between 1975 and 1990 (Jemison et al. 2006). In Prince William Sound, harbor seal counts declined by ∼63% overall between 1984 and 1997, including a 40% decline prior to the *Exxon Valdez* oil spill in 1989 (Frost et al. 1999, Ver Hoef and Frost 2003). In the Aleutian Islands, counts declined by 67% between the early 1980s and 1999, with declines of about 86% in the western Aleutians (Small et al. 2008). At Aialik Bay, a site in Kenai Fjords National Park where harbor seals haul out on glacial ice, harbor seal counts declined by 93% from 1979 to 2009 (Hoover-Miller et al. 2011). Estimated numbers of harbor seals in Johns Hopkins Inlet of Glacier Bay National Park, where the majority of seals pup and molt on ice from glaciers, declined from nearly 6,000 in 1992 to about 1,500 in 2017 (Mathews and Pendleton 2006, Womble et al. 2020). Despite the protection afforded to harbor seals in Alaska after passage of the U.S. Marine Mammal Protection Act of 1972, these examples indicated widespread declines, rather than recovery.

These declines in harbor seal numbers, mostly prior to our monitoring period, have been associated with various causes, most of which have also been touted as possible explanations for a concomitant decline in the western population of Steller sea lions (*Eumatopias jubatus*). In both cases, the declines were the most severe in the western Aleutian Islands and less severe eastward toward the Gulf of Alaska (Small et al. 2008), suggesting a common cause (Pitcher 1990). Putative causes for declines in both species include “bottom-up” (nutritional/environmental; e.g., Anderson and Piatt 1999, Rosen and Trites 2000, Trites and Donnelly 2003, Small et al. 2003, Maschner et al. 2014) and “top-down” (predation or other mortality; e.g., Springer et al. 2003, Estes et al. 2009) mechanisms. Much more research has been devoted to explaining the western Steller sea lion decline, likely because the population is listed as endangered under the U.S. Endangered Species Act. Despite considerable debate about the relative importance of the potential causes of the declines (top-down mechanisms; e.g., DeMaster et al. 2006, Wade et al. 2007, 2009, Springer et al. 2008) or (bottom-up mechanisms; Trites and Donnelly 2003, Fritz and Hinckley 2005, Calkins et al. 2013, Fritz et al. 2019), there is no consensus on which of those mechanisms are most responsible for either species’ decline (National Research Council 2003).

During our monitoring period, oceanographic conditions in Alaskan waters have intermittently reflected unprecedented warming events with marine heatwaves impacting the North Pacific in circa 2014-2016 and again in 2019-2021. Additionally, the Eastern Bering Sea experienced an extreme warm stanza from 2017-2020 (Overland and Ballinger 2025). Ecosystem responses to these events have been noted, with declines in Prince William Sound forage fish cascading up to top-level predators such as common murres (Renner et al. 2024), Steller sea lions (McHuron et al. 2024), and humpback whales (Moran et al. 2026). Seals in the Bering Sea and Aleutian Islands showed declines in body condition (Boveng et al. 2020). It is reasonable to expect similar impacts to harbor seal population trends and, while not a focus of this study, coincidental declines in some harbor seal stocks warrant future investigation.

Changes in seal abundance at glacial fjord sites, particularly in recent decades, are likely driven by environmental, bottom-up forces related to the availability of floating glacial ice, which attracted the largest aggregations of seals we observed. Overall, these sites represent 10-15% of the statewide abundance, supporting disproportionately high numbers of pups in some areas that may serve as important source populations (Streveler 1979, Calambokidis et al. 1987, Blundell et al. 2011, Crowell 2024). Although tidewater glaciers in Alaska are naturally dynamic, advancing and retreating in response to both climatic and local fjord conditions, most are thinning and retreating (McNabb and Hock 2014, Loso et al. 2026), reducing the availability of ice for seals. Recent studies have demonstrated complex links between glacial calving dynamics; ice availability; and seal distribution, abundance and behavior (Womble et al. 2021, Kaluzienski et al. 2023). Warming conditions are projected to cause melting of up to two-thirds of Alaska’s glacier ice mass by 2100 (Arendt et al. 2002, Hugonnet et al. 2021, IPCC 2023, Rounce et al. 2023, Zekollari et al. 2025). Just prior to and during our study, some long-retreating tidewater glaciers grounded (e.g., Muir, Harriman, and Tyndall Glaciers), eliminating floating ice habitat for up to 1500 seals (Streveler 1979, Sauber and Molnia 2004). At least a dozen other former tidewater glaciers in Alaska have retreated onto land since the mid-20th century (McNabb and Hock 2014, Hugonnet et al. 2021). Other fjords with some of our highest seal counts have glaciers that are rapidly thinning or retreating (e.g., Yahtse and Guyot Glaciers in Icy Bay; South Sawyer Glacier in Tracy Arm; McBride Glacier in Glacier Bay; Arendt et al. 2002, 2008, Barclay et al. 2006, O’Neel et al. 2010, Shugar et al. 2026). Given that tidewater glaciers provide pupping and molting habitat for an important segment of Alaska’s harbor seals, reduced calving of glacial ice may compound other habitat impacts and influence statewide population dynamics.

One potential top-down force that has received less scrutiny as a cause of harbor seal declines in the 1970s–1990s is a lingering demographic effect (Pitcher 1990, Crowell 2020) from widespread harvests and population control measures that occurred intermittently from the late 19th century until the onset of federal protection in 1972 (McKnight 1973, Calkins et al. 1975, Cook and Norris 1998). Crowell (2016, 2020) used a historical ecology approach to estimate that at least one million and perhaps more than 1.3 million harbor seals were killed in Alaska between 1880 and 1971. Crowell (2020) suggested that the takes were unsustainable during peak periods of removals and that the declines documented in the 1970s–1990s may have been only the tail end in a long-term loss of the vast majority of the pre-exploitation population size. Indeed, hunters in the late 1960s reported more difficulty finding seals, evidence of overharvesting during the peak years of 1964–1967 (Cook and Norris 1998). To our knowledge, though, no specific demographic mechanism has been proposed for how a period of overharvesting would cause a continued decline in the population after the cessation of significant removals. Using an age-structured population projection to model the Tugidak Island harbor seal colony, Pitcher (1990) found that the model population began to recover immediately, albeit slowly, after federal protection in 1972, despite harvest removals of the vast majority of pups in nine successive cohorts, 1964–1972. The declines continued into the 1990s at many sites in central and western Alaska after large numbers of pups were removed, suggesting that it is unlikely that successive pup removals substantially prolonged the declines. In other regions, where harvesting at pupping areas was probably less intense, other mechanisms of regulation—both bottom-up and top-down—merit further consideration as drivers for harbor seal dynamics (Crowell 2020).

In contrast to the abundance increases in 9 of the 12 stocks during the first half of our monitoring period, none experienced net increases in the second half from 2010–2023. Prince William Sound, Dixon/Cape Decision and Clarence Strait had net decreases in abundance (negative linear trend value and 95% credible interval not spanning 0.0; see Table S2.1) during the second half, and the remaining stocks had no significant trend. Thus, there was little indication of regional coherence in trends among stocks, with the possible exception of the two southernmost stocks, Dixon/Cape Decision, and Clarence Strait. Explanations or hypotheses about possible underlying causes of these patterns were beyond the scope of this study, but we hope our results provide a basis for studies focused on exploration of ecological factors that may be associated with trends in Alaska harbor seal populations over the past three decades.

### Biological insights

Maximum rates of annual per capita increase in phocid seals (given single offspring per year; female sexual maturation at 3–5 years; generation length of ∼10 years) are typically 11.5–13%, reached only in the absence of density dependent inhibition of vital rates (Härkönen et al. 2002, Bowen et al. 2003, DFO 2010). Apparent rates of increase exceeding this theoretical limit might indicate effects of immigration (i.e., lack of demographic independence between populations) or problems with the data or analytic methods. Pearson et al. (2024) used counts and haul-out data to estimate the status relative to carrying capacity of harbor seal stocks in the waters of Washington and Oregon—though they did not account for human-caused mortality, or they considered it to be subsumed in the definition of carrying capacity. Our maximum estimates of trailing 8-year trends during the period 2003– 2023 (Figure 4) ranged from 0.8% per year for the Lynn Canal/Stephens Passage stock to 7.3% per year for the North Kodiak stock; all others were less than 5% per year, well below the theoretical limit. In the absence of significant human-caused mortality, growth rates such as these, less than half of the maximum for the species, would indicate population sizes greater than half of carrying capacity (e.g., Wade 1998). The harbor seal stocks we monitored are subject to various sources of human-caused mortality, such as bycatch in coastal fisheries or the Alaska Native subsistence harvest. Most of those sources, with the possible exception of the subsistence harvest, are poorly quantified. Therefore, we made no inference from our trend estimates about the status of Alaska stocks relative to their carrying capacities or maximum net productivity levels.

Fitted relationships with explanatory variables can provide insights into the natural history and behavior of monitored populations or serve as means of verification that assessment data are sensible (i.e., are consistent with previously documented natural history patterns). During the molt period, phocid seals spend more time out of water to raise their skin temperature, thereby facilitating tissue regeneration (Feltz and Fay 1966, Ling 1974, Boily 1995). In Figure 6, the left column of plots shows that the harbor seals fitted with bio-loggers hauled out in the highest proportions in late July–early August in the Western Islands, Southwest and central Alaska, and late August–early September in Southeast Alaska and glacial fjords. These temporal ranges are broadly similar to previously reported seasonal timings for peak counts or haul-out proportions during the annual pelage molt in Alaska harbor seals (e.g., Boveng et al. 2003, Daniel et al. 2003, Simpkins et al. 2003, Mathews and Pendleton 2006, Jemison et al. 2006), though interannual or geographical variations in dates of peak counts (e.g., Daniel et al. 2003, Jemison et al. 2006) are likely to be smoothed over by our spatial and temporal pooling of the haul-out data into regional groupings. Furthermore, molt timing varies by age and sex class (Daniel et al. 2003), but because the temporal and spatial variation in molt timing is not widely monitored, it is infeasible to model the effects of age-and sex-specific timing. We assumed that the age-sex distribution among the seals with haul-out bio-loggers (Table 1) was reasonably representative of the population.

The middle column of plots in Figure 6 shows the influence of time of day on harbor seal haul-out probability; the top three rows indicate that harbor seals fitted with bio-loggers tended to haul out on intertidal shores in the highest proportions around solar noon. The peak near midday for harbor seals hauling out on shore during summer has commonly been reported in other studies from Alaska and elsewhere (Boveng et al. 2003, Simpkins et al.

2003, Cunningham et al. 2009, Mathews et al. 2016). Notably, the time of day effect was much less pronounced in Southwest and Central Alaska than in the Western Islands and Southeast Alaska. We suspect that time from low tide in Southwest and Central Alaska has a much stronger influence than time of day, due to much larger tidal exchanges than in the other two regional groupings of haul-out data.

The bottom panel of the middle column in Figure 6 shows that harbor seals hauling out on glacial ice appeared to do so in higher proportions at night. We are not aware that an inverted and less pronounced haul-out curve for seals using glacial ice has been previously described. Other studies in which harbor seals hauled out in higher proportions at night have typically associated that behavior with sources of disturbance that are more prevalent during daytime (London et al. 2012, Bankhead et al. 2023), though it can be difficult to eliminate alternative explanations (Acevedo-Gutierrez and Cendejas-Zarelli 2011). A fuller exploration of the relationship between hour of day and haul-out probability for harbor seals in glacial fjords is beyond the scope of our effort to estimate abundance during a limited portion of the year. For our purposes, we noted that the upward-trending tails of the inverted curve for hours from solar noon fell largely outside the hours of our surveys. The range of haul-out probabilities for seals in glacial fjords during our survey hours was very similar to the probabilities at intertidal haul-out sites in the Southwest, Central, and Southeast Alaska groupings of haul-out data, wherein the glacial sites are situated. Consequently, the availability adjustments for the hours of the survey counts were likewise similar among those three groupings, despite the unexplained tendency to haul out at night.

The right column of plots in Figure 6 shows the influence of tides, expressed as hours from the nearest low tide, on the probability of an individual seal being hauled out in each of the three regional groupings of haul-out data. In Southwest, Central, and Southeast Alaska, there were clear tendencies of the bio-logged seals to haul-out around the low tide, likely reflecting greater availability of preferred haul-out sites at lower water levels in those regions. The influence of tides in the Western Islands region appeared much weaker. Because the bio-loggers for this region were deployed on seals in the Aleutian Islands, where tidal ranges are small (<2.5 m during our seal counts) relative to those in the Southwest, Central, and Southeast Alaska groupings (5.2–12.3 m) the available haul-out space in the Western Islands is relatively unaffected by tide height.

Overall, the modeled effects of covariates on haul-out probability largely matched our expectations based on previous studies and our familiarity with the morphology of the coastal zones throughout the harbor seal range in Alaska. We interpret this as supportive verification that our data include meaningful variability in some of the important factors that determine the availability of harbor seals to be detected in aerial surveys. We caution, however, that covariates such as these, which may have substantial correlations among them, can easily be overinterpreted.

### Limitations and recommendations

Over the nearly three-decade duration of our study, methodologies have evolved with technology but also alongside increased fiscal constraints and changing management priorities. One constant is that our study design used a relatively small-scale (∼10–15 km) and a consistent sampling unit with seal abundance and trend being estimated for each SSU. Our statistical methods are flexible enough to allow aggregation at varying and increasing spatial scales, which would allow for new (re)analyses based on potential new stocks, adjusted stock boundaries, and areas of conservation concern. Temporal scale, however, has been a limitation. For much of the first half of our study, logistical and fiscal constraints meant any single SSU was surveyed only every five years. In the later half, a more flexible design approach has allowed some SSUs (typically those with larger numbers of seals) to be surveyed on a 3–5 year cycle but other SSUs (typically those with less seals) only every 7–10 years. Glacier Bay is a notable exception where consistent funding and effort from the NPS has supported annual monitoring from 2004 through 2023. These realities mean our insights are limited to decadal trends with model outputs showing widened confidence limits as the time since the last survey increases (see Figure 3, North/South Kodiak and Cook Inlet/Shelikoff Strait stocks). Any shorter-term perturbations (e.g., oil spills, increased anthropogenic activities, oceanographic warming anomalies) are less likely to be captured by our efforts, and our ability to respond to local stakeholder concerns regarding acute ecosystem changes might be limited. That said, our sampling design (and comprehensive time series) is well-suited to capture population responses should an intensive effort be warranted in advance of or following a known activity or environmental change that would be expected to impact harbor seal populations in Alaska.

We recommend exploring an Integrated Population Model (IPM) approach. As demonstrated in previous studies (e.g., Boveng et al., 2018; Warlick et al., 2023), IPMs can provide comparable or superior precision for abundance and trend estimates even with lower overall survey effort. Transitioning to an IPM framework would require a significant shift in survey timing and methodology. Specifically, surveys would need to be conducted during the pupping season rather than the molting period, with observers classifying seals as either pups or non-pups. This presents two significant challenges. First, harbor seal pups lack the lanugo coat common for other phocids and are, thus, not as easily distinguishable from age-one young seals. Pups are also more challenging to detect and photograph from the air—infrared imaging or similar technology would need to be employed to ensure all pups were photographed. Lastly, harbor seal pups are precocious and able to swim and dive soon after birth. Our dataset of haul-out behavior from bio-loggers does not include dependent pups, so dedicated studies to estimate the proportion of dependent pups ashore might be required. Despite these challenges, IPM reliance on pup counts would move us toward monitoring the reproductive portion of the population directly. Given the fundamental differences in haul-out behavior between molting and pupping life-history stages, any such transition must include a multi-year calibration period. During this phase, concurrent surveys in both the pupping and molting seasons would be required to ensure the continuity and comparability of the long-term Alaska-wide dataset.

Future monitoring would benefit from increased genetic sampling and new technologies to improve our estimates of stock structure. This is particularly important for those regions with limited historical sampling and that represent disproportionately large areas (e.g., Prince William Sound, Cook Inlet/Shelikof Strait, Bristol Bay, and the Aleutian Islands). More regular deployment of bio-loggers, resulting in more current data on haul-out patterns, would ensure more reliable abundance estimates and our ability to detect changes, particularly if there is covariation in the factors that affect haul-out behavior and movement (e.g., foraging behavior) with survival and reproduction.

## Conclusions

The analytical framework presented here combines a long time series of harbor seal aerial survey count data with an availability model informed by haul-out behavior that is particularly well-suited for monitoring abundance and trend in Alaska. Importantly, our implementation enables annual estimates of abundance and trend with uncertainty, even in years with limited or no survey effort. Such flexibility is critical for long-term monitoring efforts because funding support is commonly inconsistent and encumbered by unpredictability. The scalability of our approach, and the flexibility it provides, is critical in a region as large as Alaska with its logistical constraints and geographical extent.

The framework is particularly well-suited for meeting the regulatory requirements of the MMPA and providing timely estimates of abundance and trend for co-management decisions and conservation. Stock assessments could be updated annually, even in years of lower survey effort, and uncertainty surrounding those estimates grows sensibly as the time series continues away from the most recent data. Important metrics such as PBR can also be updated in a timely manner and appropriately incorporate uncertainty. Beyond the MMPA, the scalability and ability to aggregate across stocks means harbor seal population trajectories can be meaningfully included as indicators of ecosystem health within regional Ecosystem Status Reports and other fisheries management processes in support of sustainable fisheries. We also anticipate that our results will be useful for inference about marine ecosystem dynamics over nearly three decades in the Gulf of Alaska, southern Bering Sea, and Aleutian Islands.

## Acknowledgments

This paper summarizes a phase of harbor seal monitoring in Alaska led and conducted primarily by the Marine Mammal Laboratory of NOAA’s Alaska Fisheries Science Center. Our findings, however, build upon data from extensive surveys conducted by the Alaska Department of Fish and Game and the National Park Service. We are extremely grateful for the contributions of many individuals in various important capacities: Oriana Badajos, Lisa Baraff, Catherine Bell, John Bengtson, Karen Blejwas, Gail Blundell, Kaja Brix, John Burns, Jack Cesarone, Paul Conn, Michelle Cronin, Jennifer DeGroot, Kathy Frost, Henry Geijsbeek, Rhonda Hinz, Benjamin Hou, Beth Jaime, Kara Johnson, Anita Lopez, Kerry Logan, Lloyd Lowry, Bill Lucey, Barb Mahoney, Elizabeth Mathews, Anne Hoover-Miller, Sally Mizroch, John Moran, Peter Olesiuk, Ray Outlaw, Rajit Patankar, Mike Payne, Linnea Pearson, Grey Pendleton, Alana Phillips, Luciana Santos, Christine Schmale, Dana Seagars, Melissa Senac, Bob Small, Nathan Soboleff, Una Swain, Louise Taylor, Jim Thomason, Linda Vate Brattstrom, John Wells, Robin Westlake, and Kate Wynne. We are also very grateful to the many NOAA Corps and contractor pilots and maintainers that skillfully and safely operated the aircraft required for this work in remote coastal Alaska.

This work was supported primarily by funds congressionally appropriated to the National Marine Fisheries Service (NMFS) and through NMFS to the Alaska Department of Fish and Game. Funding and in-kind support were also provided by the Bureau of Ocean Energy Management, the National Park Service (Glacier Bay National Park & Preserve and the Southeast Alaska Inventory & Monitoring Network), the U.S. Navy, and the Cooperative Institute for Climate, Ocean, & Ecosystem Studies (CICOES) under NOAA Cooperative Agreement NA20OAR4320271, Contribution No.: 2026-1556.

The research presented here was conducted under the authority of the following permits: MMPA research permits include: #843, #782-1355, #782-1676, #15126, #19309, #23858. Additionally, surveys within Glacier Bay National Park were conducted under NPS permits GLBA-2009-SCI-0006, GLBA-2010-SCI-0006, GLBA-2012-SCI-0008, GLBA-2017-SCI-0020, GLBA-2023-SCI-0005.

## Author contributions

Author order for the manuscript is alphabetical. Author contributions are described following the Contributor Role Taxonomy (CRediT; https://credit.niso.org/). P. Boveng roles include conceptualization, funding acquisition, investigation, methodology, project administration, supervision, writing - original draft, writing - review and editing. G. Brady roles include data curation, investigation, methodology, validation, writing - original draft, writing - review and editing. M. Cameron roles include funding acquisition, investigation, project administration, resources, supervision, writing - review and editing. C. Christman roles include data curation, investigation, methodology, project administration, software, supervision, validation, writing - original draft, writing - review and editing. S. Dahle roles include data curation, investigation, methodology, project administration, resources, supervision, validation, writing - original draft, writing - review and editing. L. Hiruki-Raring roles include data duration, investigation, writing - review and editing. J. Jansen roles include conceptualization, data curation, investigation, methodology, project administration, resources, software, supervision, validation, writing - original draft, writing - review and editing. S. Koslovsky roles include data curation, investigation, methodology, software, supervision, validation, writing - original draft, writing - review and editing. J. London roles include conceptualization, data curation, formal analysis, investigation, methodology, project administration, resources, software, supervision, visualization, writing - original draft, writing - review and editing. B. McClintock roles include formal analysis, investigation, methodology, software, validation, visualization, writing - original draft, writing - review and editing. R. Montgomery roles include data curation, investigation, methodology, project administration, writing - review and editing. E. Moreland roles include data curation, funding acquisition, investigation, methodology, resources, software, validation, writing - original draft, writing - review and editing. E. Richmond roles include data curation, investigation, project administration, resources, software, visualization, writing - review and editing. M. Simpkins roles include conceptualization, data curation, investigation, methodology, writing - review and editing. J. Ver Hoef roles include conceptualization, formal analysis, investigation, methodology, software, validation, visualization, writing - original draft, writing - review and editing. S. Walcott roles include data curation, investigation, validation, writing - review and editing. D. Withrow roles include conceptualization, data curation, investigation, methodology, project administration, resources, supervision, writing - review and editing. J. Womble roles include data curation, funding acquisition, investigation, methodology, project administration, resources, supervision, validation, writing - original draft, writing - review and editing. K. Yano roles include data curation, investigation, validation, writing - review and editing. H. Ziel data curation, investigation, resources, validation, writing - review and editing.

## Disclaimers & Conflicts of Interest

Reference to commercial companies, products, processes, or services by trade name, trademark, manufacturer, or otherwise, does not constitute or imply its endorsement, recommendation, or favoring by the United States Government or NOAA. All authors declare no conflicts of interest relevant to this work.

