## Supplemental Material 1 for "Abundance and trends of harbor seals (*Phoca vitulina richardii*) in Alaska, 1996–2023"

### S1. Stock delineation and detailed maps

The 12 stocks of harbor seals currently identified in Alaska (Figure S1) are (1) the Aleutian Islands stock—occurring along the entire Aleutian chain from Attu Island to Ugamak Island; (2) the Pribilof Islands stock—occurring on Saint Paul and Saint George Islands, as well as on Otter and Walrus Islands; (3) the Bristol Bay stock—ranging from Nunivak Island south to the west coast of Unimak Island and extending inland to Kvichak Bay and Iliamna Lake; (4) the North Kodiak stock—ranging from approximately Middle Cape on the west coast of Kodiak Island northeast to West Amatuli Island and south to Marmot and Spruce Islands; (5) the South Kodiak stock—ranging from Middle Cape on the west coast of Kodiak Island southwest to Chirikof Island and east along the south coast of Kodiak Island to Spruce Island, including the Trinity Islands, Tugidak Island, Sitkinak Island, Sundstrom Island, Aiaktalik Island, Geese Islands, Two Headed Island, Sitkalidak Island, Ugak Island, and Long Island; (6) the Prince William Sound stock—ranging from Elizabeth Island off the southwest tip of the Kenai Peninsula to Cape Fairweather, including Prince William Sound, the Copper River Delta, Icy Bay, and Yakutat Bay; (7) the Cook Inlet/Shelikof Strait stock—ranging from the southwest tip of Unimak Island east along the southern coast of the Alaska Peninsula to Elizabeth Island off the southwest tip of the Kenai Peninsula, including Cook Inlet, Knik Arm, and Turnagain Arm; (8) the Glacier Bay/Icy Strait stock—ranging from Cape Fairweather southeast to Column Point, extending inland to Glacier Bay, Icy Strait, and from Hanus Reef south to Tenakee Inlet; (9) the Lynn Canal/Stephens Passage stock—ranging north along the east and north coast of Admiralty Island from the north end of Kupreanof Island through Lynn Canal, including Taku Inlet, Tracy Arm, and Endicott Arm; (10) the Sitka/Chatham Strait stock—ranging from Cape Bingham south to Cape Ommaney, extending inland to Table Bay on the west side of Kuiu Island and north through Chatham Strait to Cube Point off the west coast of Admiralty Island, and as far east as Cape Bendel on the northeast tip of Kupreanof Island; (11) the Dixon/Cape Decision stock—ranging from

Cape Decision on the southeast side of Kuiu Island north to Point Barrie on Kupreanof Island and extending south from Port Protection to Cape Chacon along the west coast of Prince of Wales Island and west to Cape Muzon on Dall Island, including Coronation Island, Forrester Island, and all the islands off the west coast of Prince of Wales Island; and (12) the Clarence Strait stock—ranging along the east coast of Prince of Wales Island from Cape Chacon north through Clarence Strait to Point Baker and along the east coast of Mitkof and Kupreanof Islands north to Bay Point, including Ernest Sound, Behm Canal, and Pearse Canal.

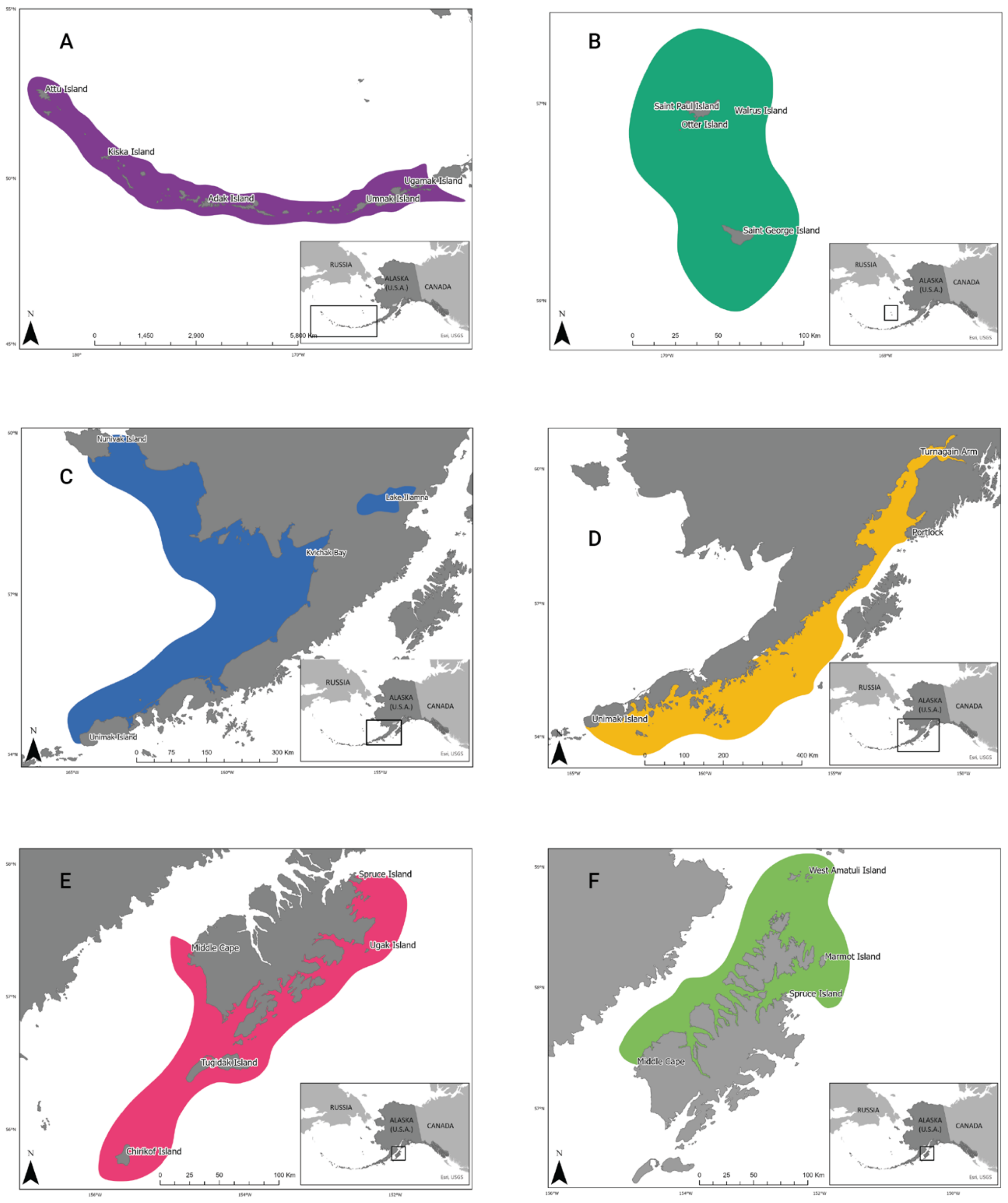


Figure S1.1. Stocks of harbor seals in Alaska: (A) Aleutian Islands, (B) Pribilof Islands, (C) Bristol Bay, (D) Cook Inlet/Shelikof Strait, (E) South Kodiak, (F) North Kodiak.

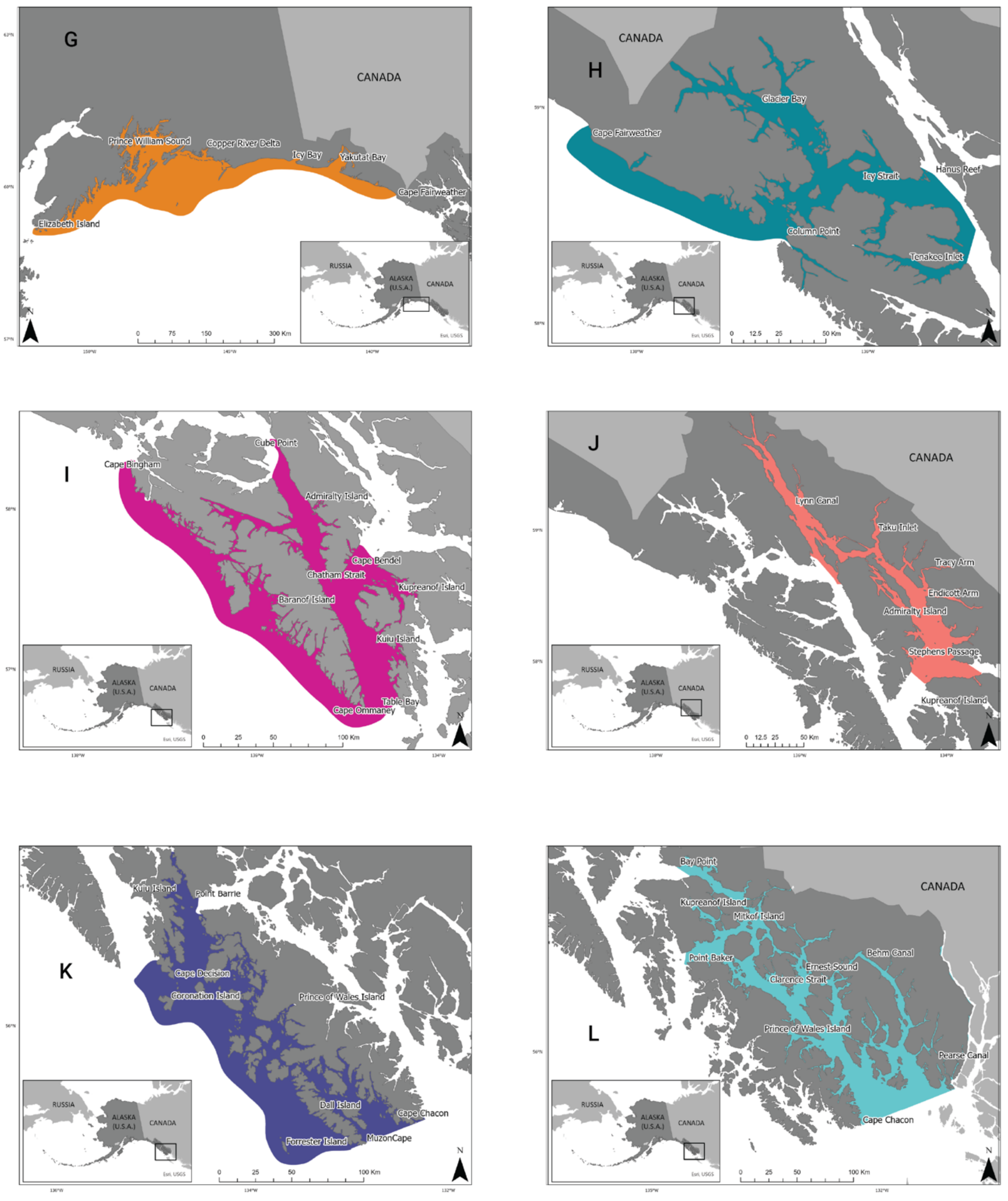


Figure S1.2 (continued). Stocks of harbor seals in Alaska: (G) Prince William Sound, (H) Glacier Bay/Icy Strait, (I) Sitka/Chatham Strait, (J) Lynn Canal/Stephens Passage, (K) Dixon/Cape Decision, and (L) Clarence Strait.
