## Supplemental Material 2 for "Abundance and trends of harbor seals (*Phoca vitulina richardii*) in Alaska, 1996–2023"

### S2. Overall Trends by Stock

| Stock | Estimate | Lower CI | Upper CI | 1st Half Est. | 1st Half Lower CI | 1st Half Upper CI | 2nd Half Est. | 2nd Half Lower CI | 2nd Half Upper CI | 1st Half Trend +/- | 1st Half Significant | 2nd Half Trend +/- | 2nd Half Significant |
|---|---|---|---|---|---|---|---|---|---|---|---|---|---|
| ALIS | 35.46 | 6.06 | 63.31 | 70.79 | 5.97 | 130.90 | -26.13 | -85.78 | 34.45 | positive | yes | negative | no |
| PRIS | -1.03 | -9.07 | 4.53 | -0.10 | -14.27 | 9.04 | -0.75 | -11.55 | 8.73 | negative | no | negative | no |
| BRBA | 274.47 | 150.61 | 391.67 | 486.33 | 227.26 | 735.32 | 249.23 | -54.53 | 555.04 | positive | yes | positive | no |
| CISH | 248.50 | 148.79 | 346.13 | 690.09 | 527.94 | 848.44 | -102.41 | -318.04 | 119.30 | positive | yes | negative | no |
| SKOD | 171.23 | 92.47 | 248.66 | 463.21 | 353.88 | 570.71 | -170.49 | -366.17 | 26.96 | positive | yes | negative | no |
| NKOD | 120.022 | 79.617 | 162.13 | 199.45 | 140.26 | 257.99 | -72.81 | -174.96 | 34.68 | positive | yes | negative | no |
| PRWS | 251.38 | 139.95 | 364.11 | 770.19 | 532.14 | 1001.65 | -926.24 | -1177.57 | -665.17 | positive | yes | negative | yes |
| GLBA | -64.52 | -108.54 | -19.40 | -274.23 | -375.96 | -187.76 | -13.81 | -102.34 | 84.76 | negative | yes | negative | no |
| SICH | 49.34 | -1.93 | 102.79 | 90.80 | 12.12 | 168.05 | -86.75 | -197.50 | 33.46 | positive | yes | negative | no |
| LCSP | -16.05 | -68.33 | 38.06 | -22.18 | -116.81 | 72.19 | -67.75 | -181.20 | 50.65 | negative | no | negative | no |
| DECD | 43.70 | -29.19 | 119.76 | 347.57 | 226.63 | 468.73 | -302.77 | -470.50 | -142.77 | positive | yes | negative | yes |
| CLST | -106.45 | -185.75 | -25.04 | 238.10 | 98.37 | 379.73 | -273.24 | -449.96 | -91.10 | positive | yes | negative | yes |
