## Supplemental Material 3 for "Abundance and trends of harbor seals (*Phoca vitulina richardii*) in Alaska, 1996–2023"

### S3 Modifications to MCMC Sampler

In order to improve mixing and better diagnose potential convergence issues, we made several key changes to the MCMC sampling for count data of Ver Hoef et al. (2025) decscribed in their Appendix A.3. For each stock, we initialized five chains with random overdispersed starting values. After adaptive tuning and burn-in, we ran each chain for as long as necessary to achieve a Gelman-Rubin-Brooks (GRB) diagnostic upper confidence interval value $< 1.2$ and an effective sample size (ESS) $> 400$ for all parameters and latent variables (e.g., Zitzmann et al., 2021). This generally required chain lengths of 1 million iterations (thinned by 1000 to save on computer storage), but there were cases where much longer chains were required (e.g. glacial sites). We were able to run much longer chains than Ver Hoef et al. (2025) because we coded the MCMC algorithm in C++ instead of R. Out of over 68,000 parameters and latent variables, there were respectively only 134 and 1 instances where these GRB or ESS criteria were not met. These instances appearred to be attributable to multimodality, but had no discernable effect on the abundance estimates across chains.

With these much longer chains that appeared to have converged, we noticed a tendency for the site-level abundance ($N_{j,t}$) estimates to drift upwards in the absence of count data (particularly when occurring at the end of a time series). We believe this is attributable to $N_{j,t}$ being bounded at zero and the random walk for $\delta_{j,t}$ being modelled on the log scale (recall that the abundance rate parameter $\lambda_{j,t} = \exp(\tau_j + \delta_{j,t})$). To discourage this tendency in the absence of data, we therefore included the weakly informative prior $\lambda_{j,t} \mid \alpha_\lambda, \beta_\lambda \sim \text{Gamma}(\alpha_\lambda, \beta_\lambda)$, with shape parameter $\alpha_\lambda = 1$ and rate parameter

$\beta_\lambda = 0.001$. We further modified the MCMC algorithm of Ver Hoef et al. (2025) to help improve mixing, as described below.

- updating $N_{j,t}$: We use essentially the same procedure as Ver Hoef et al. (2025), but use a different approach for creating the range of proposed values ($\mathbf{v}$) for $N_{j,t}$ at sites with high abundance. If there are multiple counts per year, let $v_{min}$ be the maximum count among $c_{i,j,t}; i = 1, ..., n_{j,t}$. If no counts occurred in a year, then $v_{min} = 0$. For sites with relatively low abundance, we created a range of values, $\mathbf{v} = \{v_{min}, v_{min} + 1, v_{min} + 2, \ldots, v_{max}\}$ where $v_{max}$ is the smallest integer larger than $4 \times \max(c_{i,j,t}; i = 1, 2, \ldots, n_{j,t}; t = 1996, \ldots, 2023)$ as in Ver Hoef et al. (2025). However, for sites with high abundance, $\mathbf{v}$ could be very large and sampling was slow, so we enforced that $\mathbf{v}$ have no more than 100 elements and used exponential scaling (controlled by a tuning parameter) to concentrate points around the current $N_{j,t}$ while still covering the broader interval from $v_{min}$ to $v_{max}$. If the current $N_{j,t} = v_{min}$, we used a combination of "very local" and exponential scaling for spacing the points in $\mathbf{v}$, where the first 11 elements of $\mathbf{v}$ were $\{v_{min}, v_{min} + 1, v_{min} + 2, \ldots, v_{min} + 10\}$ and the remaining elements were exponentially scaled between $v_{min} + 10$ and $v_{max}$.

- updating $\delta_{j,t}$: Proposals for $\delta_{j,t}$ were made from a uniform distribution with lower bound $l_\delta = \max(-4, \min(\delta_{j,t-1} - \omega_{j,t}, \delta_{j,t}, \delta_{j,t+1} - \omega_{j,t}))$ and upper bound $u_\delta = \min(4, \max(\delta_{j,t-1} + \omega_{j,t}, \delta_{j,t}, \delta_{j,t+1} + \omega_{j,t}))$, where $\omega_{j,t}$ is a tuning parameter. Because $\delta_{j,t}$ now influences $l_\delta$ and $u_\delta$ (and hence the proposal is no longer symmetric), the Metropolis-Hastings acceptance ratio of Ver Hoef et al. (2025) must be multiplied by $\frac{u_\delta - l_\delta}{u^*_\delta - l^*_\delta}$, where $u^*_\delta$ and $l^*_\delta$ are the upper and lower bounds evaluated at the proposed value $\delta^*_{j,t}$. Accounting for the prior on $\lambda_{j,t}$, the acceptance ratio must also be multiplied by $\frac{[\lambda|\alpha_\lambda, \beta_\lambda]}{[\lambda^*|\alpha_\lambda, \beta_\lambda]}$, where $\lambda^* = \exp\left(\tau_j + \delta^*_{j,t}\right)$.

- updating $\sigma_\delta$: Proposals for $\sigma_\delta$ were made uniformly, centered on the current value of $\sigma_\delta$, and bounded overall so that $0.00001 < \sigma_\delta < 1$. The lower and upper bounds of the uniform distribution (i.e., the proposal range around the current value of $\sigma_\delta$) were controlled by a tuning parameter. No further changes were made for updating $\sigma_\delta$.

- updating $\boldsymbol{\eta}$: Proposals for $\eta_{i,j,t}$ were made uniformly, centered on the current value of $\eta_{i,j,t}$, and bounded overall so that $-5 < \eta_{i,j,t} < 4$. The lower and upper bounds of the uniform distribution were controlled by a tuning parameter. We proposed and updated $\eta_{i,j,t}$ in blocks by site and year ($i = 1, 2, \ldots, n_{j,t}$), using the loglikelihood fragments,

$$\ell_m(\eta_{1,j,t}, \eta_{2,j,t}, \ldots, \eta_{n_{j,t},j,t} \,|\boldsymbol{\eta}_{-i,-j,-t}, \boldsymbol{\delta}, \boldsymbol{\tau}, \sigma_\delta, \boldsymbol{\gamma}, \boldsymbol{\beta}, \mathbf{c}, \mathbf{X}_c) = \sum_{i=1}^{n_{j,t}} \log(\text{Bin}(c_{i,j,t}; N_{j,t}, p_{i,j,t})) + \log(\text{N}(\eta_{i,j,t}; 0, \sigma_\eta^2)).$$

Unlike Ver Hoef et al. (2025), note that we no longer assume $\sigma_\eta^2 = 1$. We instead assume $\sigma_\eta^2$ follows an inverse gamma distribution with shape parameter $\alpha_\eta = 3$ and scale parameter $\beta_\eta = 0.5$.

- updating $\sigma_\eta^2$: Because we are no longer assuming $\sigma_\eta^2$ is fixed, we now sample it using a Gibbs step by drawing from an inverse gamma distribution with shape parameter $\alpha_\eta^* = \alpha_\eta + \frac{\sum_j \sum_t n_{j,t}}{2}$ and scale parameter $\beta_\eta^* = \beta_\eta + \frac{\sum_j \sum_t \sum_{i=1}^{n_{j,t}} \eta_{i,j,t}^2}{2}$.
